# A Promising Chemical Series of Positive Allosteric Modulators of the μ-Opioid Receptor that Enhance the Antinociceptive Efficacy of Opioids but not their Adverse Effects

**DOI:** 10.1101/2021.05.24.445057

**Authors:** Kerri D. Pryce, Hye Jin Kang, Farhana Sakloth, Yongfeng Liu, Susan Khan, Katalin Toth, Abhijeet Kapoor, Andrew Nicolais, Tao Che, Lihuai Qin, Feodora Bertherat, H. Ümit Kaniskan, Jian Jin, Michael D. Cameron, Bryan L. Roth, Venetia Zachariou, Marta Filizola

**Author notes:** These authors contributed equally to this work (Kerri D. Pryce, Hye Jin Kang, Farhana Sakloth). Department of Anesthesiology, Washington University in St. Louis School of Medicine, St. Louis, MO 63110, USA. Co-corresponding authors: Marta Filizola, PhD, Department of Pharmacological Sciences, Icahn School of Medicine at Mount Sinai, One Gustave L. Levy Place, Box 1677, New York, NY 10029, USA,; Venetia Zachariou, PhD, Nash Family Department of Neuroscience, Icahn School of Medicine at Mount Sinai, New York, NY 10029, USA One Gustave L. Levy Place, Box 1677, New York, NY 10029, USA.

## Abstract

Positive allosteric modulators (PAMs) of the µ-opioid receptor (MOR) have been proposed to exhibit therapeutic potential by maximizing the analgesic properties of clinically used opioid drugs while limiting their adverse effects or risk of overdose as a result of using lower drug doses. We herein report *in vitro* and *in vivo* characterization of two small molecules from a chemical series of MOR PAMs that exhibit: (i) MOR PAM activity and receptor subtype selectivity *in vitro*, (ii) a differential potentiation of the antinociceptive effect of oxycodone, morphine, and methadone in mouse models of pain that roughly correlates with *in vitro* activity, and (iii) a lack of potentiation of adverse effects associated with opioid administration, such as somatic withdrawal, respiratory depression, and analgesic tolerance. This series of MOR PAMs holds promise for the development of adjuncts to opioid therapy to mitigate against overdose and opioid use disorders.

## 1. Introduction

The µ-opioid receptor (MOR) subtype of the G-protein coupled receptor (GPCR) superfamily remains the main target for clinically used analgesics (Volkow and McLellan, 2016), notwithstanding both serious side effects and frequent opioid use disorder (OUD) symptoms associated with these drugs, as well as the high risk of opioid-related overdose mortality caused by activation of MORs located on brainstem neurons controlling respiration. Considering that more than 48,000 people died from overdosing on synthetic opioids other than methadone over a 12-month period ending June 2020 (data from the National Center for Health Statistics (NCHS) of the Centers for Disease Control and Prevention; https://www.cdc.gov/nchs/nvss/vsrr/drug-overdose-data.htm), safer analgesics or effective adjuncts to opioid therapy are in high demand.

Positive allosteric modulators (PAMs) of the MOR (Remesic et al., 2017) are among the medication priorities proposed by the National Institute on Drug Abuse (NIDA) to respond to the so-called “opioid epidemic” (Bruchas and Roth, 2016; Rasmussen et al., 2019; Wacker et al., 2017). The hypothesis is that MOR PAMs either enhance the activity of endogenous opioid peptides or increase the antinociceptive efficacy of clinically used opioid drugs, but not the adverse effects associated with these drugs, by virtue of allowing lower drug dosing or altering the signaling mechanisms of classical opioids.

Additional potential advantages of MOR PAMs are: (a) their expected receptor subtype selectivity due to their binding at non-conserved (and hence unique) regions on the target receptor, and (b) their possible “probe dependence” or amplification of selected downstream signaling pathways based on the specific orthosteric agonist they are used in combination with (Che and Roth, 2021; Keov et al., 2011; Wootten et al., 2012). Although some opioid receptor PAMs have also been suggested to augment functional selectivity or biased agonism towards the G-protein pathway, the exploitation of this concept for therapeutic purpose (Bohn et al., 1999; Maguma et al., 2012; Raehal et al., 2005) remains controversial (Gillis et al., 2020; Kliewer et al., 2020; Stahl and Bohn, 2020).

Notwithstanding the promise of MOR PAMs for safer and more effective use of clinically used pain medications, only a handful of MOR PAM small molecules have been published to date, and these include: BMS-986121 (Burford et al., 2013), BMS-986122 (Burford et al., 2013), BMS-986187 (Livingston and Traynor, 2018), and our very own MS1 (Bisignano et al., 2015). Unlike BMS-986122 and BMS-986187 (Livingston and Traynor, 2018), MS1 is a MOR selective PAM (Bisignano et al., 2015). While BMS-986187 and BMS-986122 have been shown to enhance G protein activation to a greater extent than β-arrestin recruitment by modulating the δ-opioid receptor (DOR) (Stanczyk et al., 2019) and MOR (Kandasamy et al., 2021), respectively, MS1 was suggested to display β-arrestin bias at MOR in the presence of endomorphin-1 or L-methadone without exhibiting activity on its own as assessed by endomorphin-1-stimulated *β*-arrestin2 recruitment assays in Chinese Hamster Ovary (CHO)-*μ* PathHunter cells (Bisignano et al., 2015). Most recently, *in vivo* experiments demonstrating that BMS-986122 produces antinociception with reduced side effects also appeared in the literature (Kandasamy et al., 2021).

Here, we report that several compounds of the MS1 chemical series exhibit PAM activity *in vitro* by increasing, to a different extent, the activity of endomorphin-1, methadone, or ([D-Ala^2^, N-MePhe^4^, Gly-ol]-enkephalin (DAMGO) in luciferase-based cAMP biosensor (GloSensor^TM^, Promega) and Tango β-arrestin recruitment assays. For two of these compounds, specifically the parent compound MS1 and the so-called compound-5 (or Comp5), we confirm (a) MOR PAM activity *in vitro* in the presence of DAMGO, methadone, morphine, or oxycodone using Bioluminescence Resonance Energy Transfer (BRET) assays, (b) MOR selectivity using radioligand binding assays and PRESTO-Tango GPCRome screening, (c) a differential enhancement of the antinociceptive effect of oxycodone, morphine, and methadone in mouse models of pain that roughly correlates with in vitro activity, and (d) a lack of potentiation of several adverse effects associated with opioid administration, such as withdrawal, analgesic tolerance, and respiratory depression.

## 2. Materials and methods

### 2.1. Selection of Chemical Compounds

The MOR PAM parent compound MS1 was used as reference in an ‘analogue by catalogue’ search of the ZINC15 database. Of the ∼100 MS1 derivatives identified, twenty-one were purchased from Princeton Biomolecular Research (Princeton, NJ) for preliminary *in vitro* screening. Additional aliquots of MS1 and Comp5 were synthesized in house for *in vivo* studies using the chemical procedures described in Supplementary Information (SI).

### 2.2. cAMP inhibition assay

To provide an initial assessment of the allosteric activity of the twenty-one selected compounds on MOR/Gα_i/o_-mediated cAMP inhibition, HEK 293T cells were co-transfected with human MOR and a luciferase-based cAMP biosensor (GloSensor^TM^, Promega). The next day, cells were seeded into Poly-lysine coated 384-well white clear bottom cell culture plates with complete DMEM + 1% dialyzed fetal bovine serum, 2 mM L-glutamine, 100 units/mL penicillin G, 100 μg/mL streptomycin at a density of 20,000 cells per 40 µL per well. The next day, the cell medium was removed and was loaded with 20 μL of assay buffer (1X HBSS, 20 mM HEPES, 0.1% BSA, pH 7.4). Then, two different concentrations (0 and 3 (1 for endomorphin-1) µM) of the tested compounds were pre-incubated for 15-20 minutes before adding 10 μL of 4X MOR agonists (endomorphin-1, DAMGO, and methadone) for another 15 minutes.

To increase the endogenous cAMP level via endogenously expressed Gs-coupled β-adrenergic receptors, 10 μL luciferin (4 mM final concentration) supplemented with isoproterenol (400 nM final concentration) were added per well. Since maximal changes in luminescence ouput were observed within 15 minutes following luciferin addition, end-point luminescence measurements were acquired at the 15 minute, using a Wallac TriLux microbeta (Perkin Elmer) luminescence counter. These luminescence measurements were used to construct dose response curves, normalized defining 0% as basal and 100% as maximal values at 0 µM concentration of the allosteric modulator, and analyzed using nonlinear regression “log(agonist) vs. response” in GraphPad Prism 9.0. Data from one experiment were collected in triplicates, and the curve fit variance (based on triplicate) was used to generate mean values of log(Emax/EC_50_) +/− standard errors.

### 2.3. β-arrestin recruitment Tango assay

The Tango assay was utilized as previously described (Kroeze et al., 2015) to provide a preliminary assessment of the allosteric activity of the 22 selected compounds on β-arrestin mediated MOR activation. Briefly, HTLA cells (which stably express β-arrestin-TEV fusion gene and tTA dependent luciferase reporter genes) were transfected with codon-optimized MOR-Tango constructs overnight. Cells were seeded into Poly-lysine coated 384-well white clear bottom cell culture plates with complete DMEM + 1% dialyzed fetal bovine serum, 2 mM L-glutamine, 100 units/mL penicillin G, 100 μg/mL streptomycin at a density of 20,000 cells per well (40 μL per well) for 6 hrs. Then, different concentrations of 10 μL of 5X tested compound prepared in the same buffer (DMEM, 1% dialyzed) were pre-incubated for 15-20 minutes before adding 10 μL of 6X MOR agonists (endomorphin-1, DAMGO, and methadone) overnight. Similar to the cAMP inhibition assay, only 2 concentrations, 0 µM and 3 µM (1 µM for endomorphin-1) of the compounds were tested. The next day, all media/drug solutions were removed and treated with 20 µL per well of BrightGlo (diluted 20-fold with Tango buffer) for 15-20 minutes followed by reading at a luminescence counter StakMax (Molecular device). These luminescence measurements were used to construct dose response curves, normalized defining 0% as the basal and 100% as maximal at 0 µM concentration of the allosteric modulator, and analyzed using nonlinear regression “log(agonist) vs. response” in GraphPad Prism 9.0. Data from one experiment were collected in triplicates, and the curve fit variance (based on triplicate) was used to generate mean values of log(Emax/EC_50_) +/− standard errors.

### 2.4. Bioluminescence Resonance Energy Transfer (BRET) assays

Comp5, which exhibited the largest EC_50_ shift between 0 μM and 3 μM in the cAMP biosensor assay with all three tested MOR orthosteric agonists, was selected and further tested in combination with methadone, DAMGO, morphine, or oxycodone using secondary G_αoA_ dissociation BRET and β-arrestin recruitment BRET assays. To measure the dissociation of labeled G_αoA_ from the labeled Gβγ complex after receptor stimulation, HEK293T cells were co-transfected in a 1:1:1:1 ratio of G_αoA_-RLuc, Gβ3, GFP2-Gγ9, and MOR construct. After 16-24 hours, transfected cells were plated in poly-l-lysine-coated 96-well white clear-bottom cell culture plates with DMEM containing 1% dialysed FBS, 100 units/mL penicillin G and 100 μg/mL streptomycin at a density of 40,000 cells in 200 μL per well and incubated overnight. The medium was aspirated and cells were washed once with 60 μL of assay buffer (1×HBSS, 20 mM HEPES, pH 7.4, 0.1% BSA). Then, 40 μL of assay buffer was loaded per well followed by addition of 20μL of 5X final concentrations (0, 0.3, 1, 3, 10 and 30 µM) of MS1 or Comp5 for 1 hour. Finally, 20 μL of 5X final concentrations of DAMGO, methadone, morphine, or oxycodone were added for another 10 min followed by addition of 20 μL of the RLuc substrate coelenterazine 400a (Nanolight), at 5 μM final concentration. Both endpoint measurements of luminescence (400 nm) and fluorescent GFP2 emission (515 nm) after 5 minutes of incubation of coelenterazine 400a were read for the plate for 1 s per well using Mithras LB940. The ratio of GFP2/RLuc was calculated per well and analyzed using a standard allosteric operational model in GraphPad Prism 9 to obtain allosteric parameters (see below for details).

To measure MOR-mediated β-arrestin2 recruitment, HEK293T cells were co-transfected in a 6:1 ratio of Venus-β-arrestin2 and MOR-RLuc. For experiments with oxycodone or morphine, GRK2 was added to increase an otherwise low signal window. After 16-24 hours, transfected cells were plated in poly-l-lysine-coated 96-well white clear-bottom cell culture plates with DMEM containing 1% dialysed FBS, 100 units/mL penicillin G and 100 μg/mL streptomycin at a density of 40,000 cells in 200 μL per well and incubated overnight. The medium was aspirated and cells were washed once with 60 μL of assay buffer (1×HBSS, 20 mM HEPES, pH 7.4, 0.1% BSA). Then, 40 μL of assay buffer were loaded per well followed by addition of 20μL of 5X final concentrations (0, 0.3, 1, 3, 10 and 30 µM) of MS1 or Comp5 for 1 hour. Finally, 20 μL of 5X drug dilution of DAMGO, morphine, oxycodone, or methadone were added for another 10 min followed by addition of 20 μL of the RLuc substrate coelenterazine h (Promega), at 5 μM final concentration. Both endpoint measurements of luminescence (485 nm) and fluorescent eYFP emission (530 nm) after 5 mins of incubation of coelenterazine h were collected for the plate for 1 s per well using Mithras LB940. The ratio of eYFP/RLuc was calculated per well and analyzed using a standard allosteric operational model in GraphPad Prism 9 to obtain allosteric parameters (see below for details).

### 2.5. Allosteric operational model and data analysis

To estimate allosteric parameters, BRET data from concentration response curves were fitted to the allosteric operational model (Christopoulos and Kenakin, 2002; Huang et al., 2015; Leach et al., 2007) in GraphPad Prism 9 as shown in the equation below:

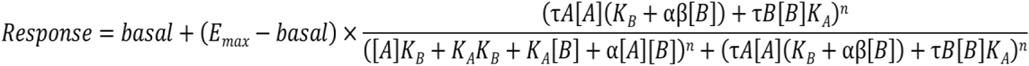

In this equation, the measured activity (Response) refers to the relative luminescence (BRET2 ratios) between the GFP2 (or eYFP) and RLuc8 emissions normalized to % orthosteric ligand response (DAMGO, morphine, oxycodone, or methadone) in the absence of allosteric compounds. Specifically, values from DAMGO, morphine, oxycodone, or methadone concentration-response curves in BRET assays in the presence of graded concentrations of MS1 or Comp5 from three independent experiments on different days were combined to generate single concentration-response curves from which to derive allosteric parameters, as well as their variance from curve fit. In the equation above, Emax corresponds to the maximal possible system response whereas ‘’basal’ is the baseline activity in the absence of test ligand. While the former is normally constrained to the maximal reading of the corresponding experiment, the latter is constrained to its baseline and set to 0 if results are normalized to fold of basal. The [A] and [B] in the equation above correspond to concentrations of the orthosteric (DAMGO, morphine, oxycodone, or methadone) and allosteric (MS1 or Comp5) ligands, respectively. In the same equation, the equilibrium dissociation constants (i.e., binding affinities) of the orthosteric agonist (DAMGO, morphine, oxycodone, or methadone) and the allosteric modulator (MS1 or Comp5) are labeled K_A_ and K_B_, respectively. Since K_B_ values of MS1 or Comp5 could not be derived directly in the MOR binding assay (Fig. S9), they were constrained to the corresponding potency values (EC_50_, the MS1 or Comp5 concentration for halfmaximal response in the GαoA BRET) in the absence of the orthosteric ligand, under the assumption that the MS1 or Comp5 potency is not significantly different from their binding affinity when the BRET assay is carried out. Since MS1 or Comp5 potency in β-arrestin2 BRET was not measurable, K_B_ on β-arrestin2 BRET was constrained to the MS1 or Comp5 potency derived in the GαoA BRET assay.

The τA and τB terms in the equation are used to indicate the ability of orthosteric and allosteric ligands to exhibit agonism, respectively. The ‘n’ in the equation is the transducer function’s slope factor that connects occupancy to response. Finally, allosteric parameters α and β correspond to the binding and efficacy cooperativity factors, respectively, providing estimates of the allosteric modulator effect on the binding affinity or efficacy of the orthosteric agonist (α/β > 1 for increased affinity/efficacy or α/β < 1 for decreased affinity/efficacy). With ‘basal’, Emax, and K_B_ constrained to their corresponding values, K_A_, τA, α, β and n are shared fitting parameters for DAMGO, morphine, oxycodone, or methadone dose response curves in the absence or presence of graded concentrations of MS1 or Comp5. For optimal comparison of the effects of MS1 and Comp5 on the different orthosteric agonists from separate estimations of α and β, and K_B_, we computed log(*αβ*/*K*_B_) values as per expert recommendation (Kenakin, 2017). The statistical significance of log(*αβ*/*K*_B_) differences at the G protein and b-arrestin channels was assessed by t-test.

### 2.6. Radioligand binding assay

Binding assays were performed using HEK293T membrane preparations transiently expressing the MOR, DOR, or KOR. Competition binding assays were set up in 96-well plates in the standard binding buffer (50 mM Tris, 0.1 mM EDTA, 10 mM MgCl_2_, 0.1% BSA, pH 7.40). Radioligand was 0.5–1 nM [^3^H]-DAMGO, [^3^H]-DADLE, or [^3^H]-U69,593 for MOR, DOR, or KOR, respectively, in standard binding buffer. Briefly, 50 μL each of [^3^H]-DAMGO, [^3^H]-DADLE, or [^3^H]-U69,593, drug solution (3X) and homogenous membrane solution (MOR, DOR, or KOR) were incubated in 96-well plate in the standard binding buffer. Reactions (either saturation or competition binding) were incubated for 2 hours at room temperature in the dark, and terminated by rapid vacuum filtration onto chilled 0.3% PEI-soaked GF/A filters followed by three quick washes with cold washing buffer (50 mM Tris HCl, pH 7.40) and read. Results (with or without normalization) were analyzed using GraphPad Prism 9.0 using one-site model.

### 2.7. GPCRome screening

To interrogate the MOR selectivity, PRESTO-Tango GPCRome screening of MS1 and Comp5 in single concentrations (10 mM) was performed at 318 GPCRs using a previously published procedure (Kroeze et al., 2015) with modifications. Briefly, HTLA cells were seeded in 384-well white plates at a density of 10000 cells/well and incubated in 40 µL of DMEM supplemented with 1% dialyzed FBS for 4 hrs. Then, the cells were transfected with 5 µL of 20 ng/well receptor Tango constructs overnight and incubated with 10 µL of 5.5 X final concentration (10 µM) compounds for another 24 hours. Media/drug solutions were removed and 20 µL/well of diluted BrightGlo were added to determine the luminescence activity. For those six receptors whose activity increased to more than 3.0-fold of basal levels of relative luminescence units, full dose–response assays were performed in the presence of MS1, Comp5, and reference (known) agonists for each receptor, except GPR151, for which no agonists are known. Thus, NECA (Tocris), Trimethylamine (Tocris), Relaxin-3 (Sigma), prostaglandin E2 (Sigma) and saquinavir (Sigma) were used as reference ligands for ADORA1, TAAR5, RXFP3, PTGER3, and BB3, respectively. Briefly, 40 µl of HTLA cells transfected with each receptor were seeded in 384-well white plates. Then the cells were loaded with 10 µl of 5 X final concentration (0, 1, 3, 10, and 30 µM) of each reference ligand as well as MS1 or Comp5 in the 1% dialyzed DMEM media for 24 hours. Results (relative luminescence units) were plotted as % of each reference ligand against individual receptors in the GraphPad 9.0 software. For GPR151, which does not have a reference ligand, results were plotted as fold of basal.

### 2.8. Animals

For behavioral experiments, we used 2- to 3-month-old male and female C57BL/6 mice (Jackson laboratories, Bar Harbor, USA). Mice were housed with a 12 h dark/light cycle with food and water ad libitum according to the Institutional Animal Care and Use Committee (IACUC) of the Icahn School of Medicine at Mount Sinai.

### 2.9. In vivo mouse pharmacokinetic (PK) studies

MS1 was taken up in 7.5% NMP, 7.5% Solutol HS, 10% propylene glycol (PEG), 15% PEG-400 and 60% 2-Hydroxypropyl-β-Cyclodextrin (HPβCD) solution (20% w/v). Comp5 was taken up in 5% NMP, 5% Solutol HS-15, 30% PEG-400 and 60% Captisol (60% w/v). A formulated solution of each compound was administered intraperitoneally (i.p.) at a 50 mg/kg dose to a designated group of 15 male Swiss Albino mice. Approximately, 60 μL of blood were acquired from three mice at each time point (0.25, 0.5, 2, 4 and 8 hours) under light isoflurane anesthesia. Blood samples were centrifuged to harvest plasma. The plasma was then stored at −70±10°C until analysis. Three mice were sacrificed immediately after blood collection to obtain brain samples for each time point at 0.25, 0.5, 2, 4 and 8 hours. Brain samples were homogenized using ice-cooled phosphate buffer saline (pH 7.4) in a ratio of 2 (buffer): 1(brain). Homogenates were stored below−70±10°C until analysis. Total homogenate volume was three times the brain weight. Pharmacokinetic analysis was conducted using the plasma and brain concentration−time data of MS1 and Comp5. The NCA module of Phoenix Win Nonlin (version 7.0) was used to perform the pharmacokinetic analysis. A fit-for-purpose LCMS/MS method was used to quantify plasma and brain samples (LLOQ for MS1 = 5.12 ng/mL for plasma and 6.15 ng/g for brain; LLOQ for Comp5 = 5.02 ng/mL for plasma and 6.03 ng/g for brain).

### 2.10. Drug formulation for in vivo mouse behavioral studies

MS1 and Comp5 were diluted in 7.5% Dimethyl sulfoxide (DMSO), 7.5% Solutol HS, 10% PEG 300, 15% PEG-400, and 60% HPβCD solution (20% w/v). Vehicle administration consist of formulation without MS1 or Comp5. Morphine sulfate, methadone hydrochloride, oxycodone hydrochloride, and naloxone hydrochloride were obtained from Sigma-Aldrich. All opioids were dissolved in sterile 0.9% saline. Formulated solutions of each compound and opioids were made fresh just before administration.

### 2.11. Hot Plate Assay

Analgesia was measured using a 52 °C hot plate apparatus (IITC Life Sciences) as previously described (Gaspari et al., 2018). Animals were habituated in the room for 30 minutes. First baseline latencies were measured, then Comp5 or MS1 was administered by i.p. injection, and 30 minutes later the opioid analgesic drug was administered by subcutaneous (s.c.) injection. Mice were tested again in the hot plate apparatus at 30 minutes after opioid treatment. A cutoff time of 40 seconds was used to avoid tissue damage and inflammation. A positive response includes jumping, licking, or flicking of either paw. For acute tolerance studies, following baseline hot plate measurements, mice were injected high opioid doses (morphine 100 mg/kg s.c. and oxycodone 80 mg/kg s.c). The next day (16 hours later) hotplate baselines were measured. Following baseline measurements, mice were injected i.p. with vehicle or Comp5 40 mg/kg or MS1 40 mg/kg and 30 min later with a low morphine (10 mg/kg s.c) or oxycodone (3.5 mg/kg s.c) dose, respectively. We assessed test latency after 30 minutes following opioid administration. Data are expressed as percentages of maximal possible effect to account for baseline differences between tested mice: [%MPE = (test latency – baseline latency)/(cutoff – baseline latency)*100].

### 2.12. CFA treatment

Left hind paw inflammation was induced by intraplantar injection of 25 μL Complete Freund’s Adjuvant ((CFA), Sigma-Aldrich, St. Louis, USA), diluted 1:1 in saline as previously described (18). CFA baseline (CFA BL) measurements were taken 18-24 hours after CFA treatment. Additional details of testing are provided in the corresponding figure legend.

### 2.13. Paw incision model of postsurgical pain

Mice were anesthetized with ketamine/xylazine and an 8-mm longitudinal incision with a no. 11 blade was made through the skin, fascia and muscle of the left plantar hind paw (Cowie and Stucky, 2019). The incision was targeted 2-mm from the proximal heel and extended toward the toes. The incision was closed with two-single sutures of 6-0 nylon (Johnson & Johnson International, New Brunswick, USA) and the paw area was treated with neomycin antibiotic ointment (Patterson Veterinary, Deven, USA). Post-surgical thermal nociceptive baseline was taken 24 hours post incision (BL after surgery) and drug treatment was tested the following day. Additional details of testing are provided in the corresponding figure legend.

### 2.14. Hargreaves Assay

Baseline thermal nociception was measured using the Hargreaves apparatus (IITC Life Sciences, California, USA) as previously described (Avrampou et al., 2019). On days 1 and 2 (habituation) mice spent 30 minutes in home cages adjusting to the testing room and were then transferred to testing chambers for 45 minutes. For thermal testing a light stimulus (IR 40) was delivered through the plexiglass floor to the plantar surface of the hind paw, and the latency to withdraw paw was measured. For each mouse, 2 measurements were taken and used to calculate the mean latency to withdrawal (seconds). A maximum IR exposure time of 15 seconds was established to ensure no tissue damage occurred, and at least 5 minutes were allowed between measurements. A positive response is considered when the experimenter observes licking, flicking, or lifting of the left hind paw. Additional details of testing are provided in the corresponding figure legends.

### 2.15. Opioid withdrawal paradigm

Mice received vehicle, Comp5 or MS1, 30 min prior to opioid or saline administration. Mice were injected every 12 hours with 10mg/kg morphine or 3.5 mg/kg oxycodone for 4 days, and on day 5 naloxone hydrochloride (1 mg/kg, s.c. Sigma-Aldrich, St. Louis, USA) was administered 1 hr after the morning opioid injection. We monitored the following withdrawal signs for 30 min after naloxone injection: i) number of jumps, wet-dog shakes, and diarrhea incidence, ii) presence of tremor and ptosis over 5 min intervals, and iii) %weight loss (weight immediately after the 30 min monitoring compared to weight immediately before naloxone administration).

### 2.16. Respiratory and Heart Rate Measurements

The impact of morphine and Comp5 on breathing and heart rates was measured in conscious/freely moving female C57Bl/6 mice using the MouseOx Plus (Starr Life Sciences Corporation). Measurements were made using neck collars sensors connected via a flexible tether with a swivel arm to allow unrestricted movement by the mouse. The hair was removed from the neck of the mice the day before the experiment. Multiplexer switching was contingent upon valid breath rate measurement for 5 seconds. Baseline measurements were continually acquired for approximately one hour before i.p. injection of morphine or Comp5 and continued for approximately one hour after dosing. Measurements were briefly interrupted when the mice were dosed. Injection time was set to time = 0. Dose levels were selected to define safety margins by comparing morphine and morphine + Comp5 doses that showed positive analgesia data to morphine doses that induce significant respiratory depression. Data represent four mice with respiratory and heart rate data combined and binned in 30-minute increments. The 25th and 75th percentiles are represented by the lower and upper error bars, respectively. Statistical tests are based on non-paired, two-sided t-tests, and standard significance labels *** p-value < 0.001, ** p-value < 0.01, n.s. p-value>0.05.

### 2.17. Statistical analysis for in vivo behavioral studies

All data are presented as mean ± SEM, with a significance set at p<0.05, denoted by the asterisk (*). Details on statistical analysis for each experiment are provided in figure legends. All data were statistically evaluated with Prism 10 software (GraphPad Software, La Jolla, California, USA). Unless otherwise indicated mice were used for a single test.

## 3. Results

### 3.1. Identification of new MOR PAMs

Using our previously discovered MOR PAM, MS1 (Bisignano et al., 2015), as reference in an ‘analogue by catalogue’ search of the ZINC15 database (Sterling and Irwin, 2015), we identified a number of derivatives of this compound, and purchased twenty-one of them for experimental testing (Supporting Information (SI) Table S1). In a search for potent MOR PAMs and expecting probe dependence (Keov et al., 2011; Wootten et al., 2012), each of these twenty-one compounds, plus the parent MS1, were preliminarily tested at two different concentrations (0 μM and 3 μM or 1 μM for endomorphin-1) in MOR/Gα_i/o_-mediated cAMP inhibition and β-arrestin recruitment Tango assays, in the presence of increasing concentrations of MOR agonists endomorphin-1, DAMGO, or L-methadone (Tables S2-S4 and Figs. S1-S6). The data revealed a very limited allosteric enhancement of G_i/o_ activity (less than 2 fold) by all tested compounds in the presence of endomorphin-1, as judged by comparing log(Emax/EC_50_) values of endomorphin-1 in MOR/Gα_i/o_-mediated cAMP inhibition assays in the presence of 0 μM and 3 μM concentrations of allosteric modulators; see Table S2. While several compounds impacted DAMGO’s and methadone’s activity more than 2 fold (compounds 2, 3, 4, 5, 7, and 8 in the case of DAMGO and compounds 5 and 6 in the case of methadone; see Table S3 and S4, respectively), Comp5 showed the largest allosteric enhancement of G_i/o_ activity by comparing EC_50_ values at 0 μM and 3 μM concentrations of allosteric modulators in the cAMP biosensor assay with all three tested MOR orthosteric agonists. Thus, this compound was selected, alongside parent compound MS1, for confirmation by G_αoA_ dissociation BRET and β-arrestin recruitment BRET assays.

Fig. 1 shows concentration-response curves of MOR obtained using G_αoA_ dissociation BRET and β-arrestin recruitment BRET assays and graded concentrations (0, 0.3, 1, 3, 10, and 30 μM) of MS1 or Comp5 in the presence of L-methadone, oxycodone, and morphine as reference orthosteric agonists. Similar concentration-response curves with DAMGO as the reference orthosteric agonist are shown in Fig. S7. Allosteric parameters were estimated based on fitting data to the allosteric operational model as described in the method section and their numerical values are shown as log(αβ), pK_B,_ and log(αβ/K_B_) in Table S5. Differences between the effect of MS1 or Comp5 on the different orthosteric agonists are shown pictorially in Fig. S8.

**Fig. 1.**
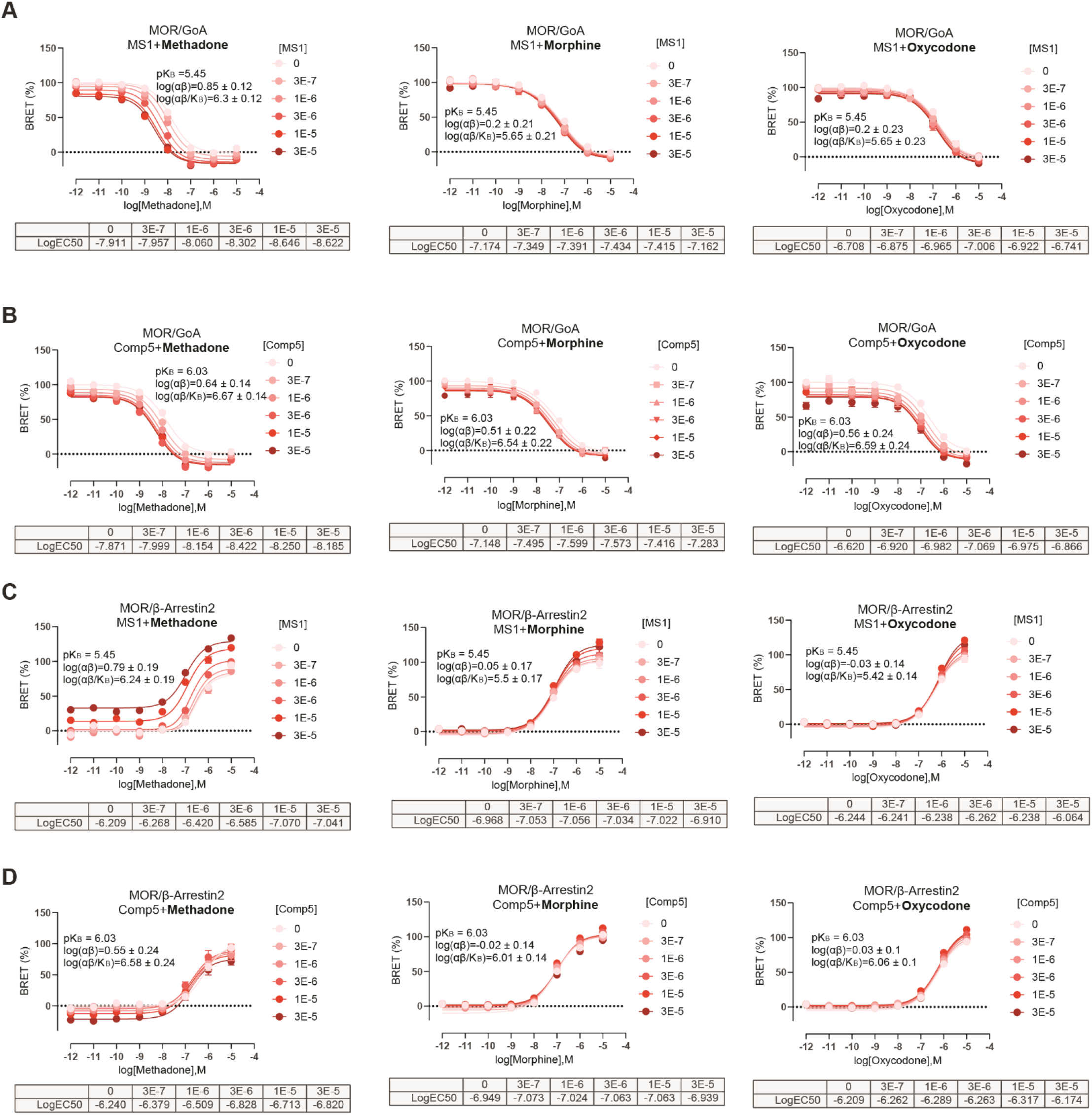
Concentration-response curves of MOR with graded concentrations of MS1 and Comp5 using L-methadone, morphine, and oxycodone as orthosteric ligands. GαoA dissociation BRET assay of MS1 (A) and Comp5 (B) and β-arrestin recruitment BRET assays of MS1 (C) and Comp5 (D) were performed to assess MS1 and Comp5 allosteric regulation of L-methadone, morphine and oxycodone, respectively, through MOR. Data are mean +/− SEM of a dataset composed of three independent experiments, each one carried out in duplicate. *K_B_* is the PAM binding affinity to free receptor, α is the binding cooperativity factor, and β is the efficacy cooperativity factor.

The larger log(αβ/K_B_) values of Comp5 or MS1 calculated for some of the orthosteric ligands (see Table S5 and Fig. S8) indicate a more significant probe dependence of MS1 compared to Comp5, with methadone being the most affected drug by MS1. Notably, these values are not significantly different when comparing each PAM effect on the GαoA or β-arrestin2 pathways, suggesting that Comp5 and MS1 exhibit a rather balanced effect on enhancing the potency of the tested orthosteric ligands based on BRET assays, with a very modest G protein-biased agonism recorded for Comp5 in combination with morphine or oxycodone, and for MS1 in combination with DAMGO.

Notably, neither MS1 nor Comp5 exhibited an effect on MOR, DOR or KOR binding (K_i_> 10 µM as per radioligand binding assays using ^3^H-DAMGO, ^3^H-DADLE ([D-Ala^2^, D-Leu^5^]-Enkephalin) and ^3^H-U69593, respectively); see Fig. S9. To assess possible off-target effects of MS1 and Comp5, PRESTO-Tango GPCRome screening (Kroeze et al., 2015) was performed, and the agonist activity of these two MOR PAMs tested at 318 non-olfactory GPCRs at 10 μM (Figs. S10A-S10B). Six receptors (ADORA1, TAAR5, RXFP3, PTGER3, BB3 and GPR151) appeared to be activated in the GPCRome screening, but the agonist activity was not observed in full dose-response assays, indicating both MS1 and Comp5 do not have off-target effects at these 6 receptors (Figs. S10C-S10H).

### 3.2. MOR PAM effect on the antinociceptive actions of clinically used opioids in mouse models of acute pain

Following a single i.p. injection of 50 mg/kg in 15 mice per compound (3 mice per time point per compound), both MS1 and Comp5 displayed good brain penetration (brain/plasma ratio of 0.96 for Comp5 and 0.75 for MS1), and maintained exposure levels over the 8-hour period post dosing (see Figs. S11A and S11B, respectively). Encouraged by the good mouse pharmacokinetic properties of these compounds, we assessed their effect on the analgesic actions of clinically used opioids (morphine, methadone, and oxycodone) in mouse models of acute nociception (exposure to noxious heat). After conducting pilot studies with low PAM doses (e.g., see Fig. S12 for Comp5) and based on the above pharmacokinetic data, a dose of 40 mg/kg i.p. was chosen for both Comp5 and MS1 for all *in vivo* experiments. Data collected using male C57Bl/6 mice in the 52°C hot plate assay show that MS1 promotes the antinociceptive action of oxycodone and methadone (Figs. 2A, 2B), but not that of morphine (Fig. 2C). In contrast, Comp5 increases antinociceptive responses to oxycodone (Fig. 2D) and morphine (Fig. 2F), but it has no effect on methadone’s action (Fig. 2E) in the hot plate assay. Using the hot plate assay, we confirmed that MS1 enhances the antinociceptive effect of methadone in female mice (Fig. S13A) and Comp5 enhances the antinociceptive effect of morphine but not methadone in female mice (Fig. S13B). We also observed that MS1 induces a small but significant extension of the antinociceptive effect of methadone in the hot plate assay (Fig. S14A (5 mg/kg s.c.)). Furthermore, Comp5-pretreated mice showed maximal antinociceptive responses for longer periods than their vehicle-treated controls when treated with high doses of oxycodone (Fig. S14B (8 mg/kg s.c.)).

**Fig. 2.**
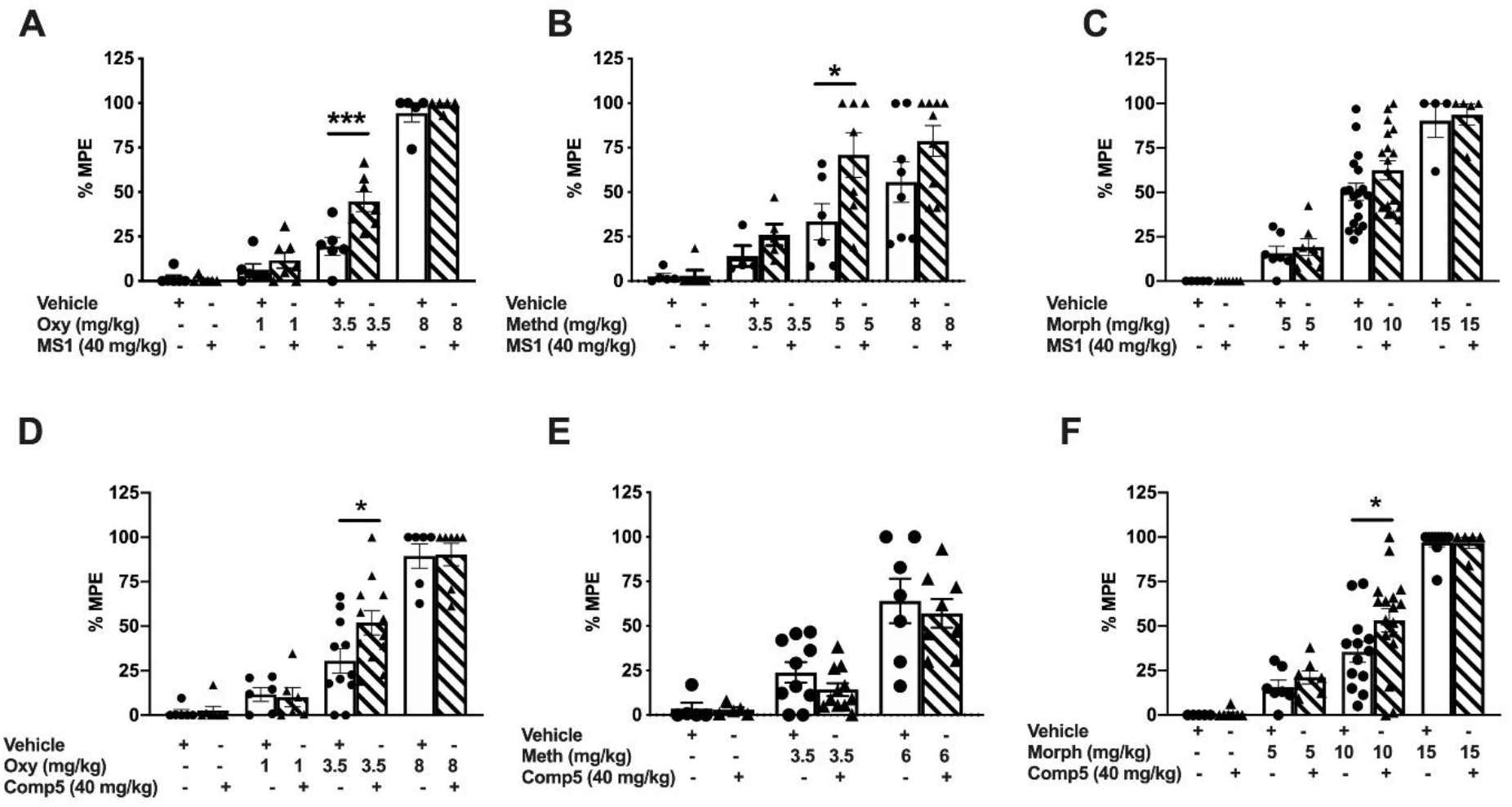
Effects of MS1 and Comp5 on antinociceptive responses to oxycodone, methadone, and morphine in the hot plate assay. Male mice were injected with MS1, Comp5, or vehicle immediately after hot plate baseline assessment, received either opioid or saline treatment 30 min later, and they were tested in the hot plate apparatus 30 min after the last injection. Data are plotted as % maximal possible effect (MPE). (A) MS1 promotes analgesic responses to oxycodone [F(7,40)=92.54, p<0.0001, (n=5-7), ***p=0.0002 for oxycodone 3.5 mg/kg +Veh vs. oxycodone 3.5 mg/kg + MS1. (B) MS1 promotes analgesic responses to methadone [F(7,42)=10.30, p<0.0001, (n=5-9 per group), *p=0.0288 for methadone 5 mg/kg +Veh vs. methadone 5 mg/kg + MS1]. (C) MS1 does not have any effect on the antinociceptive actions of morphine (n=4-18 per group). (D) Comp5 promotes the analgesic responses to oxycodone [F(9,68)= 26.52, p<0.0001, (n=6-12 per group), *p=0.0117 for oxycodone 3.5 mg/kg +Veh vs. oxycodone 3.5 mg/kg + Comp5]. (E) Comp5 has no effect on the actions of methadone (n=5-11). (F) Comp5 increases the efficacy of morphine F(7,63)= 35.43, p<0.0001, (n=5-17), *p=0.0318 for morphine 10 mg/kg +Veh vs. morphine 10 mg/kg + Comp5]. Data are reported as ± SEM. Statistics based on multiple comparisons one-way ANOVA followed by Holm-Sidak’s post hoc test. Morph (Morphine), Oxy (Oxycodone), Veh (Vehicle), and Comp5 (Compound-5). MS1, Comp5, or vehicle were injected i.p. and all opioids were injected s.c.

### 3.3. MOR PAM effect on the antinociceptive actions of opioids in mouse models of chronic pain

We next applied the CFA model of inflammatory pain to demonstrate that MS1 and Comp5 promote the antinociceptive action of opioid drugs under chronic pain states (Gaspari et al., 2018). After assessing baseline (BL) thermal hypersensitivity, mice were injected with CFA on the left hindpaw and the next day thermal hypersensitivity was verified in the Hargreaves test (CFA BL). As shown in the Fig. 3A schematic, male mice were pretreated with vehicle or Comp5 and 30 min later they were injected with either saline or morphine. Pre-treatment with Comp5 (40 mg/kg i.p.) increases Hargreaves thresholds in response to a low morphine dose (2 mg/kg s.c.) in male mice, with similar effects (at 1.5 mg/kg morphine s.c.) observed in female mice (Fig. S15A). Notably, pretreatment with Comp5 has a negligent effect on the duration of morphine antinociception (Fig. S16). Six days later, this group of male mice used for studies in Fig. 3A were tested again, in order to determine the effect of Comp5 (40 mg/kg i.p.) on antihyperalgesic responses to low oxycodone doses (0.75 mg/kg s.c.). We observed that pretreatment with Comp5 led to a significant increase in the antihyperalgesic effect of oxycodone (Fig 3B). Comparable findings were observed in sets of mice pretreated with MS1 (40 mg/kg i.p, Fig 3C). Similarly, female mice pretreated with Comp5 or MS1 (after a washout period for 6 days) show antihyperalgesic responses to morphine (1.5 mg/kg s.c.) and oxycodone (0.5 mg/kg s.c.) in the Hargreaves assay at 30 minutes after opioid treatment, whereas their vehicle-treated controls do not respond to the drug (Fig. S15A, S15B, and S15C, respectively). Parallel findings were observed in the paw incision model of post-operative pain (Kim et al., 2013) with MS1 and Comp5 promoting significant anti-hyperalgesic responses in the Hargeaves assay when combined with very low doses of methadone (2 mg/kg s.c.; Fig. 4A) or morphine (2 mg/kg s.c.; Fig. 4B), respectively.

**Fig. 3.**
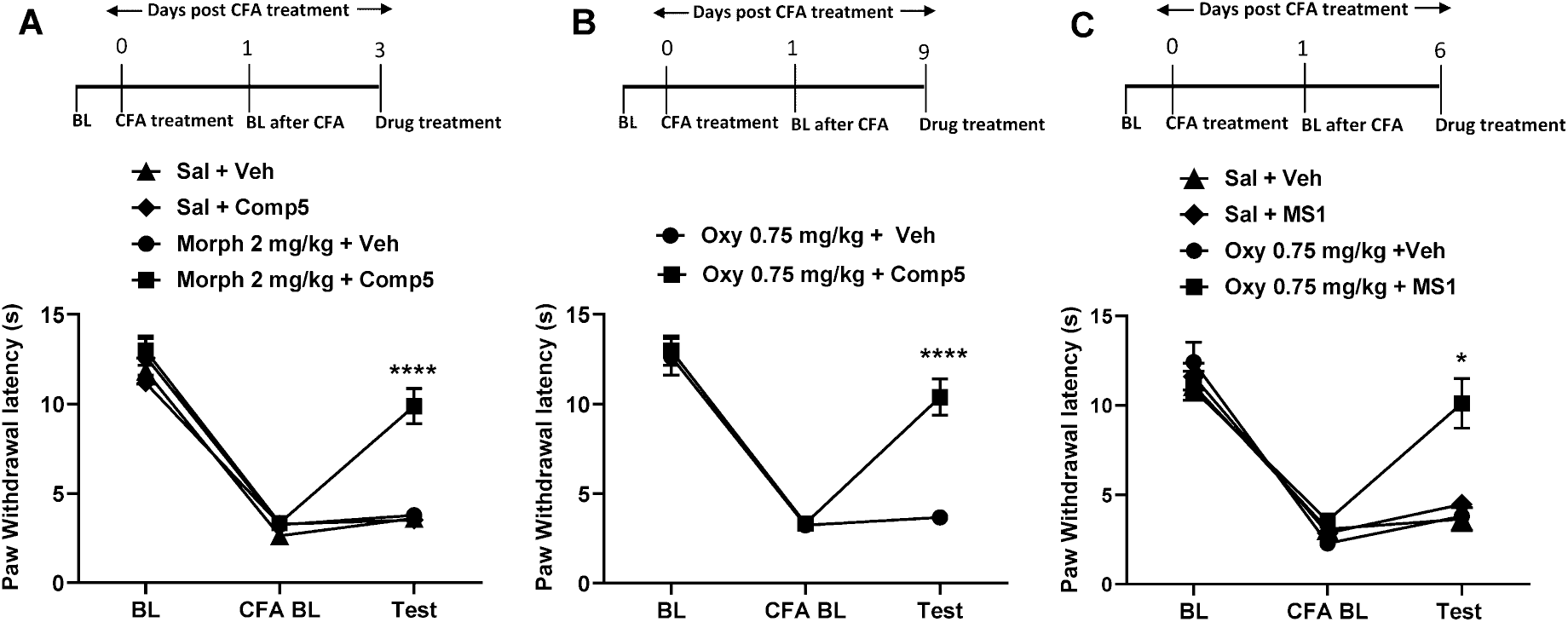
Comp5 promotes antinociceptive responses to morphine and oxycodone and MS1 increases the antinociceptive effects of morphine in a model of inflammatory pain. CFA treatment was administered to the left hindpaw after taking baseline (BL) measurements. Baseline CFA (CFA BL) measurement was taken 24 hours after CFA treatment. The mice were injected with either Comp5, MS1, or Vehicle then received either opioid treatment 30 min later, and Hargreaves responses were tested 30 min after the opioid injection. Data are reported as paw withdrawal latency in seconds. (A) In male mice, Comp5 promotes analgesic responses to morphine at 30 min [A significant interaction was reported between treatment x test time F (6,62)= 10.61, p<0.0001, (n=8 per group), ****p<0.0001 for morphine 2 mg/kg + Veh vs. morphine 2 mg/kg + Comp5, Sal+ Veh vs. morphine 2 mg/kg + Comp5, and Sal + Comp5 vs. morphine 2 mg/kg + Comp5 at 30 min]. (B) A similar effect was produced with 0.75 mg/kg Oxy [A significant interaction was reported between treatment x test time F (2,28)= 16.25, p<0.0001, (n=7-8 per group), ****p<0.0001 for Oxy 0.75 mg/kg + Veh vs. Oxy 0.75 mg/kg + Comp5]. (C) In male mice MS1 promotes analgesic responses to Oxy at 30 min [A significant interaction was reported between treatment x test time F(6,30)= 6.187, p=0.0003, (n=4-6 per group), *p<0.05 for oxycodone 0.75 mg/kg + Veh vs. oxycodone 0.75 mg/kg + MS1, Sal + Veh vs. oxycodone 0.75 mg/kg + MS1, and Sal + MS1 vs. oxycodone 0.75 mg/kg + MS1 at 30 min]. Data are reported as ± SEM. Statistics based on multiple comparisons two-way ANOVA followed by Holm—Sidak’s post hoc test. Morph (Morphine), Oxy (Oxycodone), Veh (Vehicle), and Comp5 (Compound-5). MS1, Comp5, or vehicle were injected i.p. and all opioids were injected s.c.

**Fig. 4.**
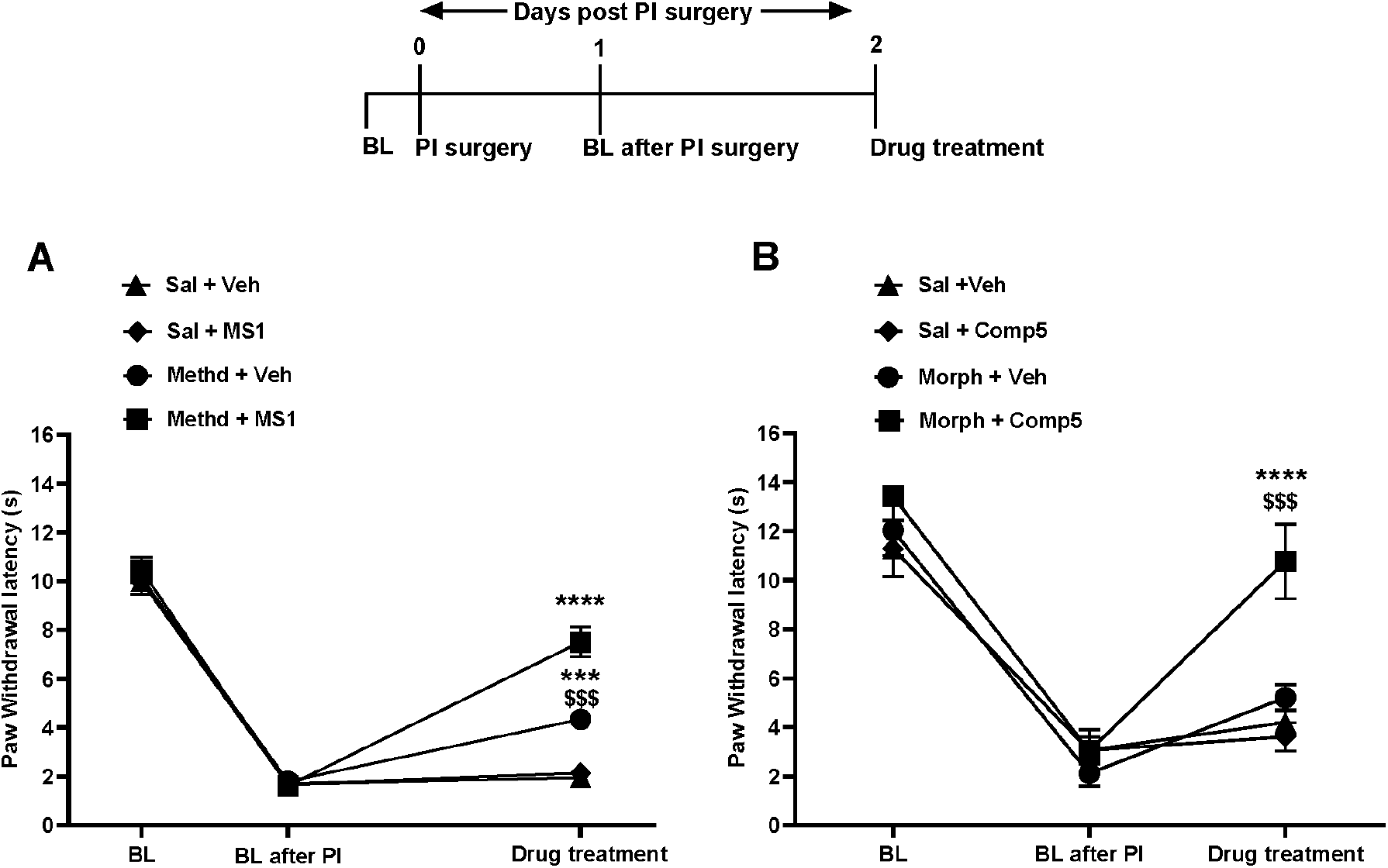
MS1 and Comp5 increase the antinociceptive effects of methadone and morphine in a model of post-operative pain. Baseline paw withdrawal latency was assessed before (BL) and after surgery. Male mice were treated with MS1 or Comp5 or vehicle 30 min before opioid administration, and Hargreaves responses were monitored 30 min after opioid treatment. (A) MS1 promotes antihyperalgesic responses to methadone [A significant interaction was reported between treatment x test time, Two-way ANOVA, F (6,46)=15.96, p<0.0001, (n=6-7), ****p<0.0001 for methadone 2 mg/kg + Veh vs. methadone 2 mg/kg + MS1, Sal + MS1 vs. methadone 2 mg/kg + MS1, Sal + Veh vs. methadone 2 mg/kg + MS1 and Sal + Veh vs. methadone 2 mg/kg + Veh, ^$$$^p<0.001 for Sal + MS1 vs. methadone 2 mg/kg + Veh]. (B) Comp5 promotes antihyperalgesic responses to morphine [A significant interaction was reported between treatment x test time, F(6,32)=5.648, p=0.0004, (n=5), ***p<0.001 for morphine 2 mg/kg + Veh vs. morphine 2 mg/kg + Comp5 and ^$$$$^p<0.0001 for Sal + Veh vs. morphine 2 mg/kg + Comp5 and Sal + Comp5 vs. morphine2 mg/kg + Comp5]. Data are reported as ± SEM. Statistics are based on multiple comparisons two-way ANOVA followed by Holm-Sidak’s post hoc test. Morph (Morphine), Methd (Methadone), Veh (Vehicle), and Comp5 (Compound-5). MS1, Comp5, or vehicle were injected i.p. and all opioids were injected s.c.

### 3.4. MOR PAM effect on opioid withdrawal signs

Prolonged opioid treatment is known to promote physical dependence, and the expression of opioid withdrawal upon discontinuation of the drug (Koob et al., 1992; Maldonado et al., 1992; Rasmussen et al., 1990). We used a naloxone-precipitated withdrawal paradigm to assess the effect of Comp5 and MS1 on the intensity of withdrawal behaviors caused by the low doses of morphine and/or oxycodone that we had used in the hot plate analgesia assay. A schematic outline of this withdrawal paradigm is shown in Fig 5A. As shown in Figs. 5B-5G, mice treated with morphine (10mg/kg s.c. twice a day) or oxycodone (3.5 mg/kg s.c. twice a day) for 5 consecutive days show a mild withdrawal syndrome. In particular, these treatment regiments lead to jumps, wet dog shakes and tremor, but we did not observe significant diarrhea or weight loss. Co-administration of Comp5 with morphine or oxycodone or MS1 with oxycodone did not affect the intensity of naloxone-precipitated withdrawal signs. PAM pretreatment did not worsen withdrawal signs. In fact, pretreatment with MS1 or Comp5 significantly reduced the incidence of wet dog shakes with oxycodone and morphine, respectively (Fig 5C and 5G). Overall, the PAMs did not produce withdrawal signs when administered without opioids (Fig 5B-5G).

**Fig. 5.**
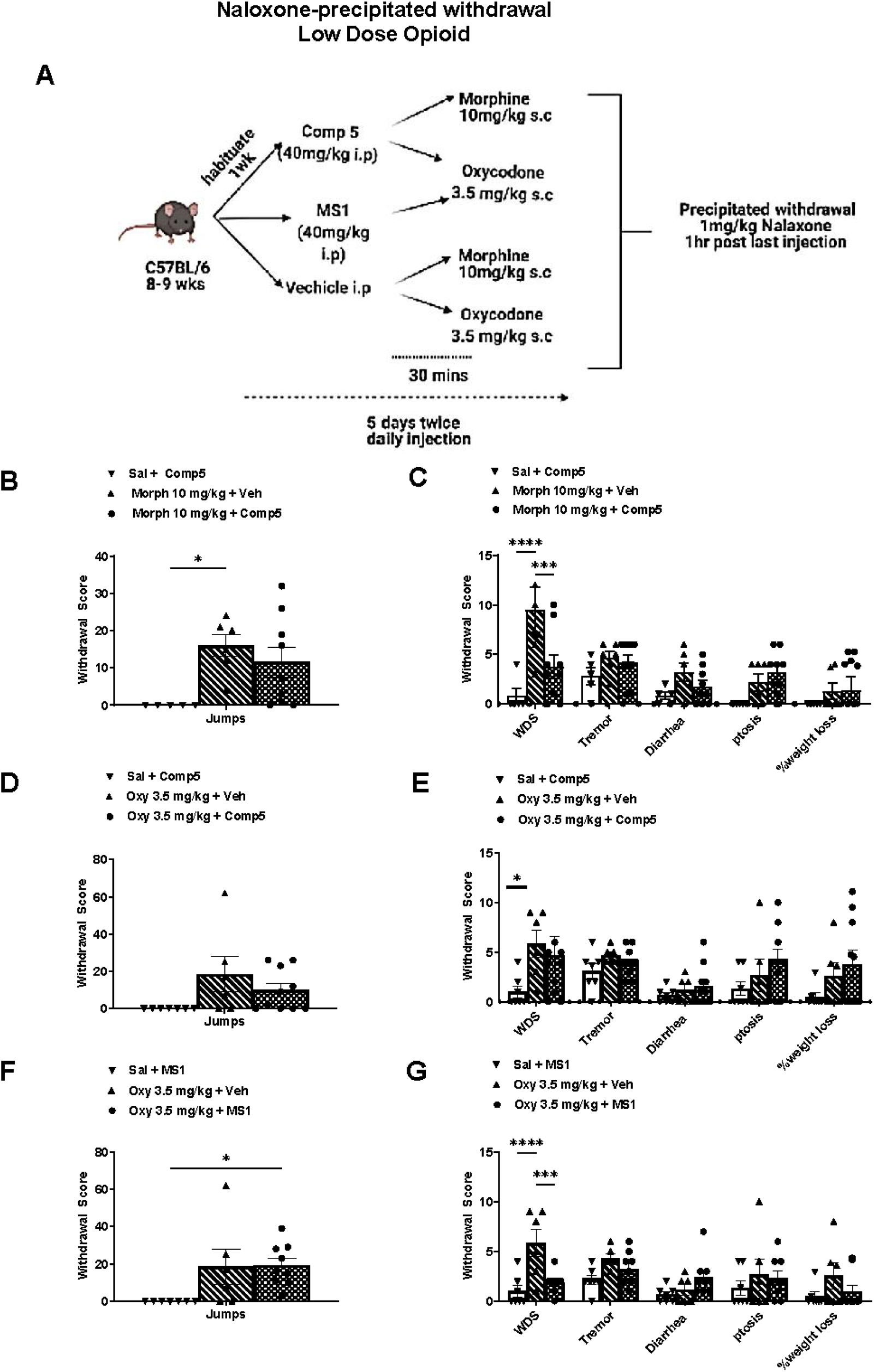
MOR PAMs do not potentiate withdrawal signs. (A) Schematic timeline for naloxone-precipitated withdrawal with low dose opioids. Comp5 or MS1 did not potentiate naloxone-precipitated morphine and/or oxycodone withdrawal symptoms (Jumps) ((B), (D), (F); 6-9 per group). (C), (G) Comp5 or MS1 pretreatment reduces wet dog shakes in naloxone-precipitated morphine and oxycodone withdrawal, respectively [Interaction F(10, 102) =2.557, p=0.0085, **** p<0.0001, *** p<0.001, ** p<0.01, *p<0.05. (C), (E), (G) Comp5 and MS1 did not affect the intensity of other symptoms (jumps, tremor, diarrhea, ptosis, and weight loss) (n=6-9 per group). Data are reported as ± SEM. Statistics based on multiple comparisons two-way ANOVA followed by Holm-Sidak’s post hoc test. Morph (Morphine), Oxy (Oxycodone) and Comp5 (Compound-5). MS1, Comp5, or vehicle were injected i.p. and all opioids were injected s.c.

### 3.5. MOR PAM effect on opioid tolerance

We investigated if Comp5 affects acute opioid tolerance using a two-day acute tolerance paradigm, in which mice were treated with high doses of morphine (100 mg/kg s.c.) or oxycodone (80 mg/kg s.c.) or with saline. The next day (16 hrs later), baseline measurements were obtained, and following that, mice were treated with vehicle, MS1 or Comp 5 (40 mg/kg i.p.) and 30 min later with a low dose of morphine (10 mg/kg s.c.) or oxycodone (3.5 mg/kg s.c.) whose analgesic response had been potentiated by the PAM. Schematic timeline of the acute tolerance paradigm is depicted in Fig 6A. Baseline hot plate latencies on test day were not different between groups (saline/morphine+vehicle=9.0±.0.6,saline/morphine+Comp5=8.89±0.34,morphine/morphine+vehicle=9.17±0.13,morphine/morphine+Comp5=8.57±0.34; Fig. 6B) or (saline/ oxycodone+vehicle=7.75±.0.42,saline/oxycodone+MS1=7.9±0.35,oxycodone/oxycodone+vehicle=8±0.7, oxycodone/oxycodone+MS1=8.8±0.7 Fig. 6C). As shown in Fig. 6B, Comp5 pretreatment leads to similar analgesic responses to those observed in vehicle groups in mice previously exposed to a high morphine dose (100 mg/kg s.c), implying no significant effect on acute analgesic tolerance to morphine in the hot plate assay. MS1 also showed no significant effect on acute analgesic tolerance to oxycodone in the hot plate assay, compared to vehicle treated controls (Fig. 6C).

**Fig. 6.**
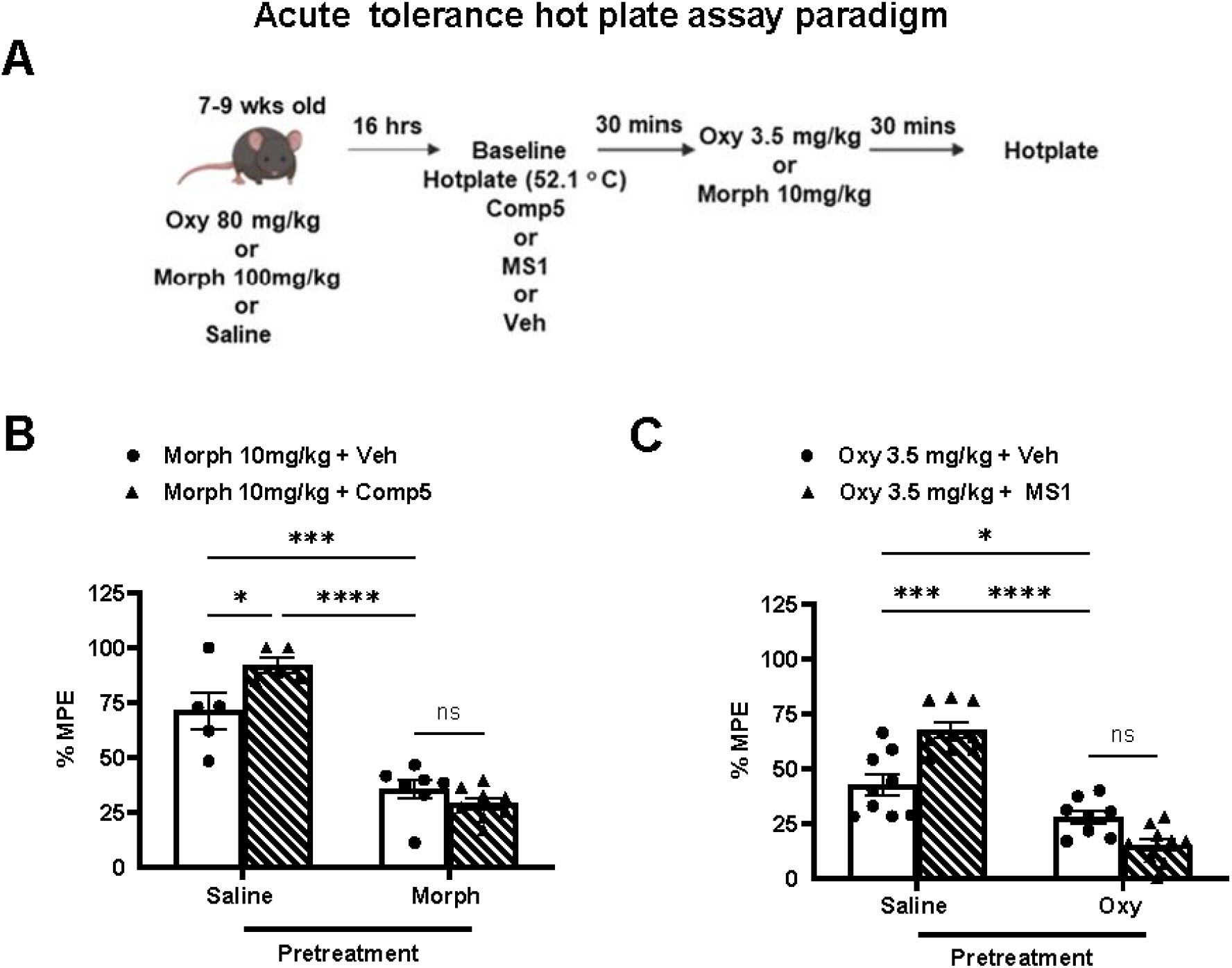
Comp5 does not affect tolerance. (A) Schematic timeline of acute tolerance paradigm. Mice were treated with high dose morphine (100 mg/kg s.c.) or high dose oxycodone (80 mg/kg s.c.) or saline; 16 hrs later baseline hotplate plate latencies are assessed. Mice were then injected with Comp5, or MS1, or vehicle immediately after hot plate baseline assessment and 30 min later injected with 10 mg/kg s.c. morphine or 3.5 mg/kg s.c. oxycodone, respectively. Data are plotted as % MPE. (B) Comp5 shows no significant effect on acute analgesic tolerance to morphine in the hot plate assay (F(3, 30)= 12.32, p<0.0001, ****p<0.0001, *** p<0.001, *p<0.05 (n= 6-9 per group)). (C) Likewise, MS1 demonstrate no significant effect on acute analgesic tolerance to oxycodone in the hot plate assay (F(3, 31)= 17.32, p<0.0001, ****p<0.0001, *** p<0.001, *p<0.05 (n=6-9 per group)). MS1, Comp5, or vehicle were injected i.p. and all opioids were injected s.c.

### 3.6. MOR PAM effect on morphine-induced respiratory depression

We evaluated the impact of Comp5 on another common on-target side effect of opioid drugs, that is, respiratory depression. Specifically, we measured breathing and heart rates in conscious/freely moving female C57Bl/6 mice using the apparatus and procedure described in Methods. As shown in Figure 7, compared with vehicle, a high dose of morphine (30 mg/kg s.c.) caused significant respiratory depression, accompanied by a decrease in heart rates. In contrast, by itself or in combination with the very low dose of morphine (2mg/kg s.c.) whose antinociceptive response is capable of potentiating, Comp5 (40 mg/kg i.p.) did not have a significant effect on either breathing or heart rates.

**Figure 7.**
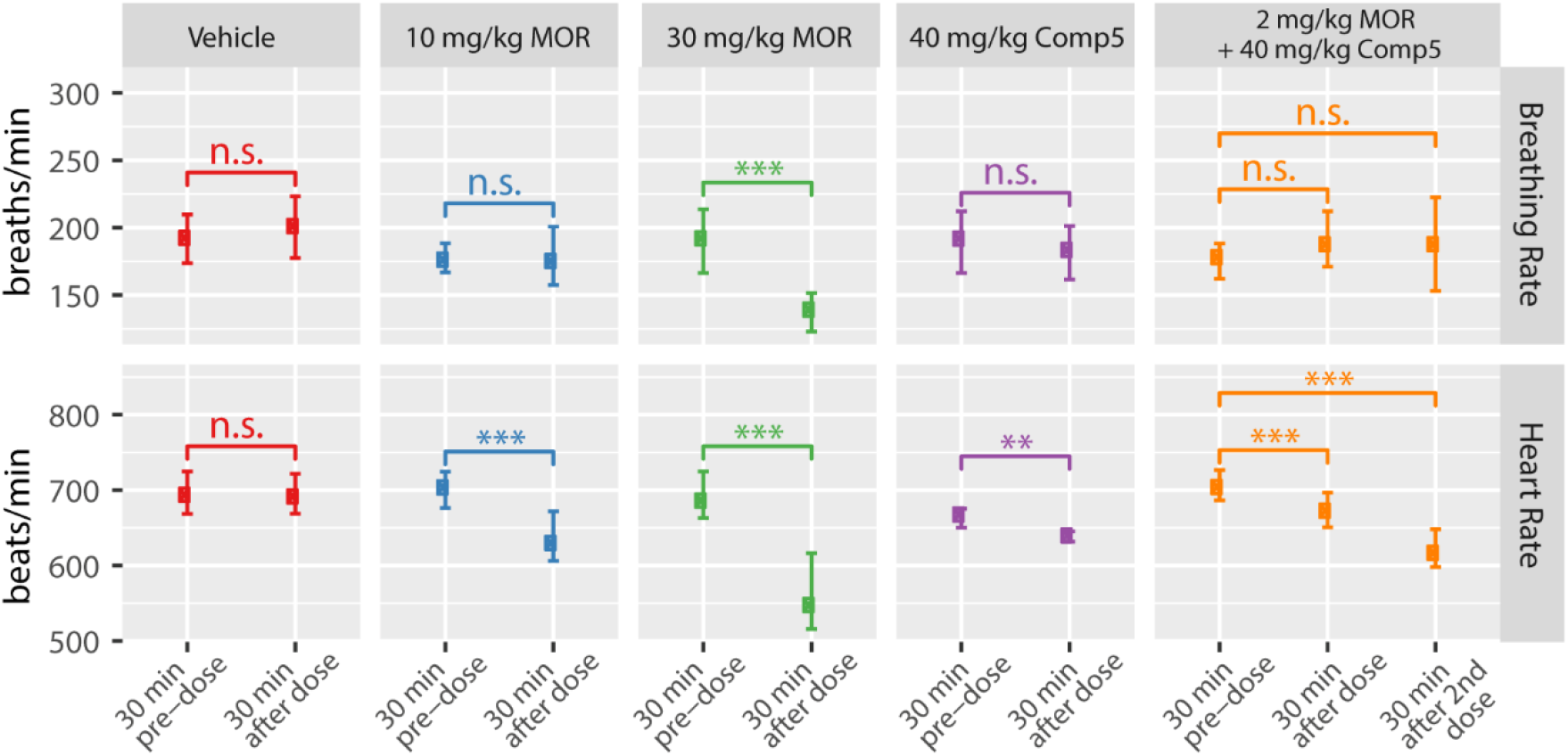
Comp5 has no impact on breathing and heart rates. Breathing and heart rates were measured in conscious/freely moving female C57Bl/6 mice using the MouseOx Plus apparatus. Data represent four mice with respiratory and heart rate data combined and binned in 30-minute increments. The lower and upper error bars represent the 25th and 75th percentiles, respectively. Statistics are based on non-paired, two-sided t-tests. For breathing rates, T = −1.11, 0.16, 7.68, and 0.75, for Vehicle, 10 mg/kg MOR, 30 mg/kg MOR, and 40 mg/kg Comp5, respectively. For 2 mg/kg MOR + 40 mg/kg Comp5, T=−1.01 and −0.97 for comparison of the 30-min pre-dose with the 30 min after the first dose and with the 30 min after the second dose; *** p < 0.001, ** p < 0.01, n.s. p>0.05; number of observations=12-38 per group. For heart rates, T=0.21, 5.76, 8.15, 3.14, for Vehicle, 10 mg/kg MOR, 30 mg/kg MOR, and 40 mg/kg Comp5, respectively. For 2 mg/kg MOR + 40 mg/kg Comp5, T=3.81 and 8.62 for comparison of the 30-min pre-dose with the 30 min after the first dose and with the 30 min after the second dose; *** p < 0.001, ** p< 0.01, n.s. p >0.05; number of observations=10-36 per group.

## 4. Discussion

Although still considered the gold-standard for the treatment of severe pain, opioid analgesics are fraught with a number of adverse effects that preclude their effective use. Finding ways to make these drugs safer rather than embarking in costly drug discovery campaigns aimed at identifying novel analgesics targeting opioid receptors and/or working through different targets and largely unexplored neurochemical mechanisms, constitutes a desirable alternative. One of the proposed ways to make opioid drugs safer to use is by limiting their dosing via allosteric modulation. Specifically, MOR PAMs would boost the antinociceptive effects with minimal dosing of the opioid drugs and consequent reduction of their adverse effects. However, this hypothesis has yet to be fully demonstrated *in vivo* with only one study (Kandasamy et al., 2021), published while our work was under review, reporting potentiation of the analgesic response of morphine, but not its adverse effects, by a single MOR PAM.

A notable difference between this recently published study (Kandasamy et al., 2021) and the work reported herein is that our compounds do not exert a significant PAM effect on the endogenous opioid peptide endomorphin-1, but rather their effects are more pronounced on a number of exogenous MOR agonists. While we decided to focus on Comp5, the MS1 derivative that produced the largest activity shift with all three tested MOR agonists (endomorphin-1, L-methadone and DAMGO) in the primary cAMP biosensor assay, it is important to note that there are several other compounds of the MS1 series that may be worthy of follow-up studies based on their observed PAM activity in the G protein or β-arrestin channels. Another advantage of our study is that we verified potentiation of antinociception for more than one compound and opioid combination, in acute noxious heat pain, inflammatory, and post-operative pain models. Notably, we were able to observe significant antinociceptive effects at very low opioid doses in the presence of Comp5 or MS1, compared to the doses used to suppress thermal hyperalgesia in models of peripheral inflammation (Armendariz and Nazarian, 2018; Gaspari et al., 2018). Similarly, using the plantar incision assay, a model of post-operative pain whose induced thermal hypersensitivity can be treated with 3 mg/kg of morphine (Cowie and Stucky, 2019), we could observe antinociception with the PAMs at reduced doses of opioids (2 mg/kg) which by themselves would produce negligible antinociception.

As expected and seen in other systems, the MOR PAMs reported here showed a clear probe dependence, which was not always the same between *in vitro* and *in vivo* experiments. In line with BRET data, Comp5 was found to potentiate the antinociceptive response to morphine and oxycodone in both male and female mice in different pain models. However, at odds with the *in vitro* data, Comp5 did not potentiate the antinociceptive response to methadone in the hotplate assay. Notably, while the BRET-inferred allosteric modulation of methadone, oxycodone, morphine, and DAMGO by Comp5 and MS1 is comparable, our *in vitro* data support a more significant probe dependence of MS1 as compared to Comp5. Although additional experiments would be required with negative drug combinations to unambiguously establish whether the ligand-dependence is consistent across assays, the lack of correspondence between data from *in vitro* and *in vivo* experiments is neither surprising nor unprecedented given the limitations of both experimental settings.

Our data also revealed a lack of correlation between signaling bias (as inferred from BRET assays) and specific responses to opioid drugs. This observation is in line with recent evidence (Bachmutsky et al., 2021; Gillis et al., 2020; Kliewer et al., 2020) that challenges the concept of opioid analgesic effects being chiefly regulated by G protein signaling while adverse effects may be mediated by β-arrestin (see (Che et al., 2021) for a recent review). Although Comp5 and MS1 exhibit a very modest G protein-biased agonism when combined with morphine/oxycodone and DAMGO, respectively, they mostly perform as balanced ligands in the BRET assay. Notably, the conclusions from BRET assays on MS1 are at odds with our early findings based on endomorphin-1-stimulated *β*-arrestin2 recruitment assays in CHO-*μ* PathHunter cells (Bisignano et al., 2015), which suggested that MS1 acted as a β-arrestin-biased MOR PAM by virtue of enhancing the potency of L-methadone in recruiting β-arrestin more than activating G-protein. However, we believe the bias quantification from BRET-based assays reported herein may be more accurate as it is based on more direct measurements and a superior quantification strategy. Future work will be required to assess PAM activity at other Gα subunit pathways, thus enabling a better understanding of the observed differences between *in vitro* and *in vivo* effects of the tested PAMs.

Notwithstanding their identity of balanced ligands, Comp5 and MS1 both exhibited promising side effect profiles in that they did not exacerbate the expression of somatic withdrawal symptoms or respiratory depression and did not promote analgesic tolerance, further contributing to the hypothesis that MOR PAMs may widen the opioid therapeutic window. Notably, we report that morphine-induced respiratory depression is observed at doses three to five times what is required for analgesia. However, in the presence of the MOR PAM Comp5, analgesia is achieved at lower doses, tripling the respiratory safety index. While more studies are obviously required to determine whether MOR PAMs can enable a safer use of opioids in humans, the studies reported here represent a step forward towards this goal.

In summary, our findings suggest that the MS1 series represents a promising lead series worthy of further investigation for the development of effective adjuncts to opioid therapy. Among the advantages of this chemical series with respect to other series published to date, are (a) a verified suitability for structure-activity relationship studies based on the demonstrated MOR PAM activity of several commercially available MS1 derivatives, (b) a demonstrated selectivity for MOR, (c) an inability to exert antinociception by themselves notwithstanding some recorded PAM activity *in vitro* in the presence of low concentrations of the endogenous *MOR* agonist endomorphin-1, but not verified *in vivo* (i.e., Comp5 and MS1 have no effect on their own), and (d) a chemical scaffold whose synthesis is modular and amenable to rapid modifications to multiple regions, making it ideally suited for future drug development studies.

## Credit authorship contribution statement

**Kerri Pryce**: Investigation, Formal analysis, Visualization, Writing – review and editing. **Hye Jin Kang**: Investigation, Formal analysis, Visualization, Writing – review and editing. **Farhana Sakloth**: Investigation, Formal analysis, Visualization, Writing – review and editing. **Yongfeng Liu**: Investigation, Formal analysis, Writing – review and editing. **Susan Khan:** Investigation, Formal analysis. **Katalin Toth:** Investigation, Formal analysis. **Abhijeet Kapoor**: Investigation, Visualization. **Andrew Nicolais**: Investigation, Formal analysis. **Tao Che**: Investigation, Formal analysis, Visualization, Writing – review and editing. **Lihuai Qin**: Investigation. **Feodora Bertherat**: Investigation. **H. Ümit Kaniskan**: Supervision, Writing – review and editing. **Bryan L. Roth**: Supervision, Methodology, Resources, Funding acquisition, Writing – review & editing. **Jian Jin**: Supervision, Resources, Writing – review & editing. **Venetia Zachariou**: Conceptualization, Supervision, Methodology, Resources, Funding acquisition, Writing – review & editing. **Marta Filizola**: Conceptualization, Supervision, Methodology, Resources, Funding acquisition, Writing – original draft, review and editing.

## Acknowledgements

The authors would like to thank Dr. Terry Kenakin for his recommendations on the quantification of signaling bias and allosteric parameters, as well as Dr. Davide Provasi for producing Figure 7. This work was supported by National Institutes of Health grants to M.F. (DA045473), V.Z. (NS086444, NS111351, and DA08227), and B.L.R. (271201800023C-P00001-9999-1). Computations were run on resources available through the Office of Research Infrastructure of the National Institutes of Health under award numbers S10OD018522 and S10OD026880.

## ABBREVIATIONS

ADORA1: adenosine A1 receptor
BB3: bombesin-like receptor 3
BRET: bioluminescence resonance energy transfer
BSA: bovine serum albumin
CFA: Complete Freud’s Adjuvant
CFA BL: Complete Freud’s Adjuvant baseline
Comp5: compound-5
DAMGO: [D-Ala^2^, N-MePhe^4^, Gly-ol]-enkephalin
DMEM: Dulbecco’s modified eagle medium
DMSO: dimethyl sulfoxide
DOR: δ-opioid receptor
FBS: fetal bovine serum
GPCR: G-protein coupled receptor
HBSS: Hank’s balanced salt solution
KOR: κ-opioid receptor
MOR: µ-opioid receptor
NIDA: National Institute on Drug Abuse
OUD: opioid use disorder
PAM: positive allosteric modulator
PRESTO: parallel receptorome expression and screening via transcriptional output
PTGER3: prostaglandin EP3 receptor
RXFP3: relaxin family peptide receptor 3
TAAR5: trace amine-associated receptor 5

