## Supplemental Material for "A Promising Chemical Series of Positive Allosteric Modulators of the μ-Opioid Receptor that Enhance the Antinociceptive Efficacy of Opioids but not their Adverse Effects"

[1]*Nash Family Department of Neuroscience, Icahn School of Medicine at Mount Sinai, New York, NY 10029, USA;* [2]*Department of Pharmacology, University of North Carolina at Chapel Hill School of Medicine, Chapel Hill, NC 27599, USA;* [3]*Department of Molecular Medicine, The Scripps Research Institute, Jupiter, FL 33458, USA;* [4]*Department of Pharmacological Sciences, Icahn School of Medicine at Mount Sinai, New York, NY 10029, USA;* [5]*Mount Sinai Center for Therapeutics Discovery, Department of Oncological Sciences, Tisch Cancer Institute, Icahn School of Medicine at Mount Sinai, New York, NY 10029, USA*

Materials and Methods for Synthetic Chemistry……………………………………………pg. 3

Table S1…………………………………………………………………………..…pg. 6

Table S2…………………………………………………………………………..…pg. 9

Table S3……………………………………………………………..………………pg. 10

Table S4………………………………..………………………………………………pg. 11

Table S5………………………………..………………………………………………pg. 12

Scheme 1……………………………..…………………………………………………pg. 13

Fig. S1……………………………………………………………………….……….pg. 14

Fig. S2……………………………………………………………………….……….pg. 15

Fig. S3……………………………………………………………………….……….pg. 16

Fig. S4……………………………………………………………………….………pg. 17

Fig. S5……………………………………………………………………………..………pg. 18

Fig. S6……………………………………………………………………………..………pg. 19

Fig. S7……………………………………………………………………………..………pg. 20

Fig. S8……………………………………………………………………………..………pg. 21

Fig. S9……………………………………………………………………………..………pg. 22

Fig. S10……………………………………………………………………………..………pg. 23

Fig. S11……………………………………………………………………………..………pg. 24

Fig. S12……………………………………………………………………………..………pg. 25

Fig. S13……………………………………………………………………………..………pg. 26

Fig. S14……………………………………………………………………………..………pg. 27

Fig. S15……………………………………………………………………………..………pg. 28

Fig. S16……………………………………………………………………………..………pg. 29

### Chemistry General Procedures

All chemical reagents were purchased from commercial vendors and used for *in vitro* studies without further purification. The flash column chromatography was conducted using a Teledyne ISCO CombiFlash Rf$^+$ instrument. This instrument was also equipped with a variable-wavelength UV detector and a fraction collector. RediSep Rf normal phase silica columns were used for purification. High-performance liquid chromatography (HPLC) spectra for compounds were acquired using an Agilent 1200 Series system with a DAD detector. Chromatography was performed on a 2.1 × 150 mm Zorbax 300SB-$C_{18}$ 5 μm column with water containing 0.1% formic acid as solvent A and acetonitrile containing 0.1% formic acid as solvent B at a flow rate of 0.4 mL/min. The gradient program was as follows: 1% B (0−1 min), 1−99% B (1−4 min), and 99% B (4−8 min). Ultraperformance liquid chromatography (UPLC) spectra for compounds were acquired using a Waters Acquity I-Class UPLC system with a PDA detector. Chromatography was performed on a 2.1 × 30 mm ACQUITY UPLC BEH $C_{18}$ 1.7 μm column with water containing 3% acetonitrile, 0.1% formic acid as solvent A and acetonitrile containing 0.1% formic acid as solvent B at a flow rate of 0.8 mL/min. The gradient program was as follows: 1−99% B (1−1.5 min), and 99−1% B (1.5−2.5 min). High-resolution mass spectra (HRMS) data were acquired in the positive ion mode using Agilent G1969A API-TOF with an electrospray ionization (ESI) source. Nuclear magnetic resonance (NMR) spectra were acquired on either a Bruker DRX500 spectrometer (500 MHz $^1$H) or a Bruker Avance-III 800 MHz spectrometer (201 MHz $^{13}$C). Chemical shifts are reported in ppm (δ).

#### Methyl ((4-bromophenyl)sulfonyl)glycinate (1)

To a solution of glycine methyl ester hydrochloride (250 mg, 2 mmol) and triethylamine (0.7mL, 4.8mmol) in dichloromethane was added 4-Bromobenzenesulfonyl chloride (561 mg, 2.2 mmol) at 0 °C. The resulting solution was warmed to room temperature (rt) and stirred for 1h. The reaction was washed with brine. The organic phase was dried over sodium sulfate and concentrated. The resulting residue was purified by silica gel column to yield the title compound as white solid (410 mg, 67%). $^1$H NMR (800 MHz, DMSO-$d_6$) δ

8.36 (t, *J* = 6.2 Hz, 1H), 7.81 (d, *J* = 8.1 Hz, 2H), 7.73 (d, *J* = 8.1 Hz, 2H), 3.75 (d, *J* = 6.1 Hz, 2H), 3.54 (s, 3H). MS *m/z* $[M + H]^+$ calcd. for $C_9H_{11}BrNO_4S^+$ 308.0, 310.0; found 308.2, 310.0.

**Methyl N-benzyl-N-((4-bromophenyl)sulfonyl)glycinate (2)**

Sodium hydride (80 mg, 2 mmol, 60% in mineral oil) was added in portions to a solution of **1** (410 mg, 1.3 mmol) in dry dimethylformamide (3 mL) at ice bath. The mixture was stirred for 30 min at the same temperature until evolution of hydrogen was ceased, then benzyl bromide (178 μL, 1.4 mmol) was added. The resultant mixture was stirred for 1 h at room temperature. After cooling with ice bath, water was added slowly to quench the excess of sodium hydride. The phases were then separated and the aqueous phase was extracted with ethyl acetate. Combined organic phases was washed with water and brine, dried over anhydrous sodium sulfate and concentrated. The residue was purified by silica gel column to yield the title compound as yellow solid (390 mg, 74%). $^1H$ NMR (800 MHz, DMSO-$d_6$) δ 7.85 – 7.79 (m, 4H), 7.35 – 7.29 (m, 3H), 7.25 (d, *J* = 7.4 Hz, 2H), 4.42 (s, 2H), 3.99 (s, 2H), 3.47 (s, 3H). MS *m/z* $[M + H]^+$ calcd. for $C_{16}H_{17}BrNO_4S^+$ 398.0, 400.0; found 398.0, 400.0.

***N*-benzyl-*N*-((4-bromophenyl)sulfonyl)glycine (3)**

A solution of **2** (390 mg, 1mmol) in THF, $H_2O$, MeOH was added anhydrous LiOH (28mg, 1.2 mmol) to. The solution was stirred at rt overnight. Then mixture was concentrated and 1N HCl was added to adjust the pH to 3. After extraction with ethyl acetate, the organic solvent was combined. Combined organic phases was washed with water and brine, dried over anhydrous sodium sulfate and concentrated. The residue was purified by silica gel column to yield the title compound as white solid (320 mg, 85%). $^1H$ NMR (500 MHz, DMSO-$d_6$) δ 7.83 – 7.77 (m, 4H), 7.36 – 7.27 (m, 3H), 7.26 – 7.22 (m, 2H), 4.44 (s, 2H), 3.86 (s, 2H). MS m/z $[M - H]^-$ calcd. for $C_{15}H_{13}BrNO_4S^-$ 382.0, 384.0; found 382.0, 384.0.

**2-((*N*-benzyl-4-bromophenyl)sulfonamido)-*N*-(3-chloro-4-methoxyphenyl)acetamide (MS1)**

3-Chloro-4-methoxyaniline (500 mg, 3.2 mmol) added to a solution of **3** (1.2 g, 3.2 mmol), EDCI (730 mg, 3.8 mmol), HOAt (517 mg, 3.8 mmol), DIEA (792 µL, 4.8 mmol) in 10 mL $CH_2Cl_2$, the resulting mixture was stirred overnight. Then the organic phase was washed with water and brine, dried over anhydrous sodium sulfate and concentrated. The residue was purified by silica gel column to yield the title compound as white solid (1.4 g, 86%). $^1$H NMR (500 MHz, DMSO-$d_6$) δ 9.91 (s, 1H), 7.84 – 7.76 (m, 4H), 7.57 (d, *J* = 2.5 Hz, 1H), 7.38 – 7.22 (m, 6H), 7.08 (d, *J* = 9.0 Hz, 1H), 4.51 (s, 2H), 3.95 (s, 2H), 3.82 (s, 3H). $^{13}$C NMR (201 MHz, DMSO) δ 166.03, 151.16, 139.37, 136.18, 132.62, 132.43, 129.69, 129.01, 128.77, 128.24, 127.22, 121.53, 121.01, 119.68, 113.35, 56.64, 51.88, 49.21. HRMS *m/z* $[M + H]^+$ calcd. for $C_{22}H_{21}BrClN_2O_4S^+$ 523.0088, 525.0068; found 523.0084, 525.0059.

**2-((*N*-benzyl-4-bromophenyl)sulfonamido)-*N*-(3-chloro-2-methylphenyl)acetamide (compound-5)**

3-Chloro-2-methylaniline (37 mg, 0.26 mmol) added to a solution of **3** (0.1 g, 0.26 mmol), EDCI (61 mg, 0.32 mmol), HOAt (44 mg, 0.32 mmol), DIEA (65 µL, 0.39 mmol) in 2 mL $CH_2Cl_2$, the resulting mixture was stirred overnight. Then the organic phase was washed with water and brine, dried over anhydrous sodium sulfate and concentrated. The residue was purified by silica gel column to yield the title compound as white solid (0.1 g, 76%). $^1$H NMR (500 MHz, DMSO-$d_6$) δ 9.50 (s, 1H), 7.88 – 7.78 (m, 4H), 7.41 – 7.26 (m, 6H), 7.20 – 7.11 (m, 2H), 4.50 (s, 2H), 4.04 (s, 2H), 2.11 (s, 3H). $^{13}$C NMR (201 MHz, DMSO) δ 166.63, 139.35, 137.56, 136.15, 134.20, 132.66, 130.79, 129.75, 129.06, 128.82, 128.31, 127.25, 126.68, 124.80, 51.94, 48.95, 15.38. HRMS m/z $[M + H]^+$ calcd for $C_{22}H_{21}BrClN_2O_3S^+$ 507.0139, 509.0119; found 507.0114, 509.0095.

**Table S1:** ZINC identification codes for MS1 (compound #18) and twenty-one derivatives.

| **Compound** | ZINC ID | 2D Structure |
|---|---|---|
| 1 | ZINC000001062898 | 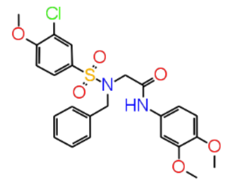 |
| 2 | ZINC000001079721 | 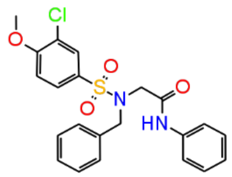 |
| 3 | ZINC000009720734 | 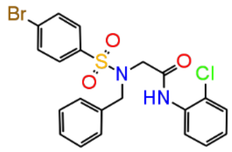 |
| 4 | ZINC000009720735 | 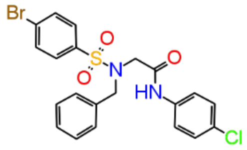 |
| 5 | ZINC000009720746 | 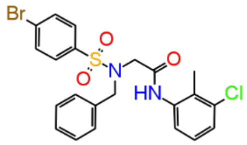 |
| 6 | ZINC000009720747 | 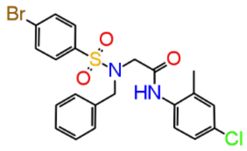 |
| 7 | ZINC000009720748 | 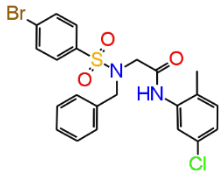 |

| | | |
|---|---|---|
| 8 | ZINC000009720755 | 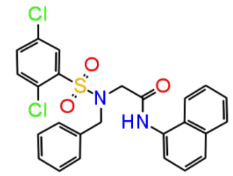 |
| 9 | ZINC000009720876 | 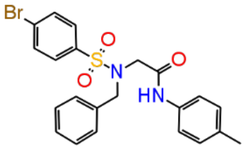 |
| 10 | ZINC000009720877 | 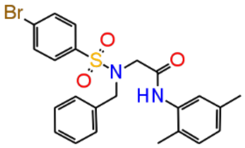 |
| 11 | ZINC000009720888 | 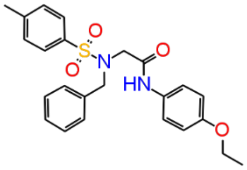 |
| 12 | ZINC000001208422 | 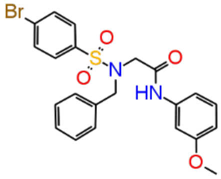 |
| 13 | ZINC000008721844 | 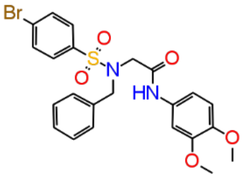 |
| 14 | ZINC000009720732 | 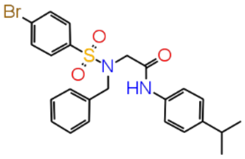 |
| 15 | ZINC000009720733 | 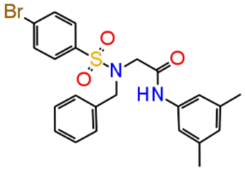 |

| | | |
|---|---|---|
| 16 | ZINC000009720749 | 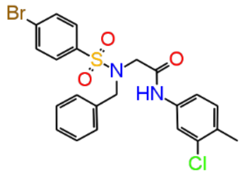 |
| 17 | ZINC000009720878 | 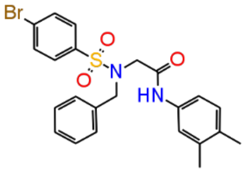 |
| 18 | ZINC000001206097 (MS1) | 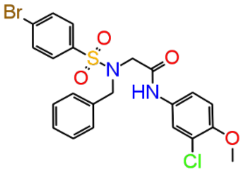 |
| 19 | ZINC000032109715 | 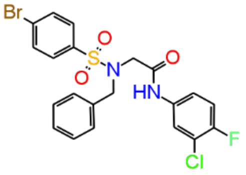 |
| 20 | ZINC000408619623 | 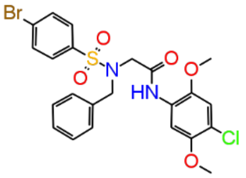 |
| 21 | ZINC000408979717 | 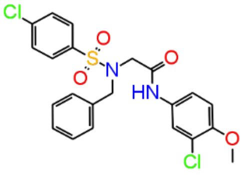 |
| 22 | ZINC000409184325 | 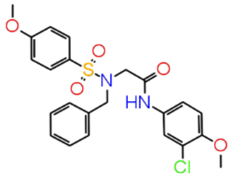 |

**Table S2:** log(Emax/EC50) values of endomorphin-1 in luciferase-based cAMP biosensor (GloSensor™, Promega) and Tango β-arrestin recruitment assays in the absence (0) or presence (1 μM) of MS1 (compound #18) and its derivatives. Data from one experiment were collected in triplicates, and the curve fit variance (based on triplicate) was used to generate mean values of log(Emax/$EC_{50}$) +/- standard errors.

| Compound | log(Emax/EC50) ± SEM | | | | log(Emax/EC50) ± SEM | | | |
|---|---|---|---|---|---|---|---|---|
| | cAMP inhibition assay | | | | Tango arrestin recruitment | | | |
| | 0 | 1E-6 (M) | delta | fold | 0 | 1E-6 (M) | delta | fold |
| 1 | 8.45 ± 0.05 | 8.39 ± 0.05 | -0.06± 0.07 | 0.9 | 5.89 ± 0.05 | 6.23 ± 0.08 | 0.34 ± 0.09 | 2.2 |
| 2 | 8.45 ± 0.06 | 8.35 ± 0.08 | -0.1± 0.1 | 0.8 | 5.56 ± 0.09 | 5.98 ± 0.06 | 0.42 ± 0.11 | 2.6 |
| 3 | 8.35 ± 0.10 | 8.44 ± 0.07 | 0.09± 0.12 | 1.2 | 5.67 ± 0.11 | 6.1 ± 0.07 | 0.43 ± 0.13 | 2.7 |
| 4 | 8.35 ± 0.10 | 8.41 ± 0.1 | 0.06± 0.14 | 1.1 | 6.01 ± 0.07 | 5.61 ± 0.11 | -0.4 ± 0.13 | 0.4 |
| 5 | 8.27 ± 0.09 | 8.43 ± 0.08 | 0.16± 0.12 | 1.4 | 5.74 ± 0.07 | 5.78 ± 0.08 | 0.04 ± 0.11 | 1.1 |
| 6 | 8.27 ± 0.09 | 8.28 ± 0.1 | 0.01± 0.13 | 1 | 5.74 ± 0.07 | 5.71 ± 0.07 | -0.03 ± 0.1 | 0.9 |
| 7 | 8.34 ± 0.13 | 8.47 ± 0.09 | 0.13± 0.16 | 1.3 | 5.69 ± 0.11 | 5.52 ± 0.11 | -0.17 ± 0.16 | 0.7 |
| 8 | 8.34 ± 0.13 | 8.36 ± 0.13 | 0.02± 0.18 | 1 | 6.01 ± 0.08 | 5.95 ± 0.1 | -0.06 ± 0.13 | 0.9 |
| 9 | 8.24 ± 0.10 | 8.37 ± 0.08 | 0.13± 0.13 | 1.3 | 5.95 ± 0.07 | 5.91 ± 0.09 | -0.04 ± 0.11 | 0.9 |
| 10 | 8.25 ± 0.09 | 8.24 ± 0.1 | -0.01± 0.13 | 1 | 5.95 ± 0.07 | 5.99 ± 0.1 | 0.04 ± 0.12 | 1.1 |
| 11 | 8.15 ± 0.08 | 8.25 ± 0.05 | 0.1± 0.09 | 1.3 | 5.50 ± 0.12 | 6.17 ± 0.11 | 0.67 ± 0.16 | 4.7 |
| 12 | 8.15 ± 0.08 | 8.06 ± 0.1 | -0.09± 0.13 | 0.8 | 5.50 ± 0.12 | 5.63 ± 0.1 | 0.13 ± 0.16 | 1.3 |
| 13 | 8.26 ± 0.13 | 8.34 ± 0.08 | 0.08± 0.15 | 1.2 | 5.74 ± 0.12 | 5.62 ± 0.12 | -0.12 ± 0.17 | 0.8 |
| 14 | 8.26 ± 0.13 | 8.15 ± 0.15 | -0.11± 0.2 | 0.8 | 5.75 ± 0.12 | 5.72 ± 0.12 | -0.03 ± 0.17 | 0.9 |
| 15 | 8.25 ± 0.09 | 8.3 ± 0.07 | 0.05± 0.11 | 1.1 | 6.16 ± 0.08 | 5.91 ± 0.08 | -0.25 ± 0.11 | 0.6 |
| 16 | 8.25 ± 0.08 | 8.18 ± 0.09 | -0.07± 0.12 | 0.9 | 6.16 ± 0.08 | 5.96 ± 0.11 | -0.2 ± 0.14 | 0.6 |
| 17 | 8.28 ± 0.11 | 8.28 ± 0.08 | 0± 0.14 | 1 | 5.87 ± 0.08 | 6.1 ± 0.08 | 0.23 ± 0.11 | 1.7 |
| 18 | 8.25 ± 0.09 | 8.11 ± 0.09 | -0.14± 0.13 | 0.7 | 5.87 ± 0.08 | 5.8 ± 0.07 | -0.07 ± 0.11 | 0.9 |
| 19 | 8.26 ± 0.08 | 8.39 ± 0.05 | 0.13± 0.09 | 1.3 | 6.04 ± 0.13 | 5.55 ± 0.14 | -0.49 ± 0.19 | 0.3 |
| 20 | 8.23 ± 0.09 | 8.12 ± 0.09 | -0.11± 0.13 | 0.8 | 6.04 ± 0.13 | 5.65 ± 0.08 | -0.39 ± 0.15 | 0.4 |
| 21 | 8.05 ± 0.08 | 8.28 ± 0.07 | 0.23± 0.11 | 1.7 | 5.99 ± 0.08 | 6.16 ± 0.08 | 0.17 ± 0.11 | 1.5 |
| 22 | 8.05 ± 0.08 | 8.02 ± 0.08 | -0.03± 0.11 | 0.9 | 5.99 ± 0.08 | 5.82 ± 0.13 | -0.17 ± 0.15 | 0.7 |

**Table S3:** log(Emax/EC50) values of DAMGO in luciferase-based cAMP biosensor (GloSensor$^{TM}$, Promega) and Tango β-arrestin recruitment assays in the absence (0) or presence (3 μM) of MS1 (compound #18) and its derivatives. Data from one experiment were collected in triplicates, and the curve fit variance (based on triplicate) was used to generate mean values of log(Emax/$EC_{50}$) +/- standard errors.

| Compound | log(Emax/EC50) ± SEM<br>cAMP inhibition assay | | | | log(Emax/EC50) ± SEM<br>Tango arrestin recruitment | | | |
|---|---|---|---|---|---|---|---|---|
| | 0 | 3E-6 (M) | delta | fold | 0 | 3E-6 (M) | delta | fold |
| 1 | 8.30 ± 0.06 | 8.38 ± 0.05 | 0.08 ± 0.08 | 1.2 | 6.38 ± 0.11 | 6.61 ± 0.09 | 0.23 ± 0.14 | 1.7 |
| 2 | 8.30 ± 0.06 | 8.65 ± 0.04 | 0.35 ± 0.07 | 2.2 | 6.07 ± 0.08 | 6.16 ± 0.07 | 0.09 ± 0.11 | 1.2 |
| 3 | 8.33 ± 0.06 | 8.66 ± 0.04 | 0.33 ± 0.07 | 2.1 | 5.87 ± 0.06 | 6.27 ± 0.06 | 0.4 ± 0.08 | 2.5 |
| 4 | 8.33 ± 0.06 | 8.78 ± 0.1 | 0.45 ± 0.12 | 2.8 | 6.07 ± 0.05 | 6.51 ± 0.09 | 0.44 ± 0.1 | 2.8 |
| 5 | 8.37 ± 0.05 | 8.73 ± 0.04 | 0.36 ± 0.06 | 2.3 | 6.07 ± 0.06 | 6.15 ± 0.09 | 0.08 ± 0.11 | 1.2 |
| 6 | 8.37 ± 0.05 | 8.67 ± 0.05 | 0.3 ± 0.07 | 2 | 6.30 ± 0.09 | 6.7 ± 0.04 | 0.4 ± 0.1 | 2.5 |
| 7 | 8.26 ± 0.05 | 8.59 ± 0.05 | 0.33 ± 0.07 | 2.1 | 5.67 ± 0.07 | 6.45 ± 0.06 | 0.78 ± 0.09 | 6 |
| 8 | 8.26 ± 0.05 | 8.6 ± 0.04 | 0.34 ± 0.06 | 2.2 | 6.07 ± 0.10 | 6.86 ± 0.04 | 0.79 ± 0.11 | 6.2 |
| 9 | 8.35 ± 0.05 | 8.57 ± 0.04 | 0.22 ± 0.06 | 1.7 | 5.79 ± 0.05 | 6.12 ± 0.07 | 0.33 ± 0.09 | 2.1 |
| 10 | 8.35 ± 0.05 | 8.52 ± 0.05 | 0.17 ± 0.07 | 1.5 | 6.00 ± 0.08 | 6.09 ± 0.07 | 0.09 ± 0.11 | 1.2 |
| 11 | 8.26 ± 0.04 | 8.31 ± 0.04 | 0.05 ± 0.06 | 1.1 | 5.68 ± 0.11 | 6.33 ± 0.05 | 0.65 ± 0.12 | 4.5 |
| 12 | 8.26 ± 0.05 | 8.49 ± 0.04 | 0.23 ± 0.06 | 1.7 | 5.73 ± 0.11 | 6.01 ± 0.12 | 0.28 ± 0.16 | 1.9 |
| 13 | 8.33 ± 0.05 | 8.34 ± 0.04 | 0.01 ± 0.06 | 1 | 5.78 ± 0.06 | 6.21 ± 0.03 | 0.43 ± 0.07 | 2.7 |
| 14 | 8.33 ± 0.05 | 8.56 ± 0.05 | 0.23 ± 0.07 | 1.7 | 6.04 ± 0.09 | 6.16 ± 0.07 | 0.12 ± 0.11 | 1.3 |
| 15 | 8.27 ± 0.04 | 8.27 ± 0.04 | 0 ± 0.06 | 1 | 5.92 ± 0.03 | 5.77 ± 0.05 | -0.15 ± 0.06 | 0.7 |
| 16 | 8.27 ± 0.04 | 8.58 ± 0.04 | 0.31 ± 0.06 | 2 | 5.90 ± 0.05 | 6.18 ± 0.06 | 0.28 ± 0.08 | 1.9 |
| 17 | 8.32 ± 0.05 | 8.37 ± 0.04 | 0.05 ± 0.06 | 1.1 | 5.93 ± 0.07 | 6.03 ± 0.06 | 0.1 ± 0.09 | 1.3 |
| 18 | 8.32 ± 0.05 | 8.59 ± 0.02 | 0.27 ± 0.05 | 1.9 | 6.06 ± 0.07 | 6.79 ± 0.07 | 0.73 ± 0.1 | 5.4 |
| 19 | 8.15 ± 0.05 | 8.26 ± 0.05 | 0.11 ± 0.07 | 1.3 | 5.95 ± 0.04 | 6.03 ± 0.11 | 0.08 ± 0.12 | 1.2 |
| 20 | 8.15 ± 0.05 | 8.35 ± 0.04 | 0.2 ± 0.06 | 1.6 | 6.01 ± 0.06 | 6.65 ± 0.05 | 0.64 ± 0.08 | 4.4 |
| 21 | 8.23 ± 0.07 | 8.39 ± 0.06 | 0.16 ± 0.09 | 1.4 | 6.11 ± 0.11 | 6.7 ± 0.06 | 0.59 ± 0.13 | 3.9 |
| 22 | 8.23 ± 0.07 | 8.32 ± 0.04 | 0.09 ± 0.08 | 1.2 | 6.05 ± 0.08 | 6.13 ± 0.04 | 0.08 ± 0.09 | 1.2 |

**Table S4:** log(Emax/EC50) values of methadone in luciferase-based cAMP biosensor (GloSensor™, Promega) and Tango β-arrestin recruitment assays in the absence (0) or presence (3 μM) of MS1 (compound #18) and its derivatives. Data from one experiment were collected in triplicates, and the curve fit variance (based on triplicate) was used to generate mean values of log(Emax/$EC_{50}$) +/- standard errors.

| Compound | log(Emax/EC50) ± SEM | | | | log(Emax/EC50) ± SEM | | | |
|---|---|---|---|---|---|---|---|---|
| | cAMP inhibition assay | | | | Tango arrestin recruitment | | | |
| | 0 | 3E-6 (M) | Delta | fold | 0 | 3E-6 (M) | Delta | fold |
| 1 | 7.71 ± 0.07 | 7.77 ± 0.06 | 0.06 ± 0.09 | 1.1 | 5.68 ± 0.05 | 6.12 ± 0.04 | 0.44 ± 0.06 | 2.8 |
| 2 | 7.82 ± 0.08 | 7.85 ± 0.08 | 0.03 ± 0.11 | 1.1 | 6.05 ± 0.05 | 6.34 ± 0.05 | 0.29 ± 0.07 | 1.9 |
| 3 | 7.72 ± 0.13 | 7.94 ± 0.11 | 0.22 ± 0.17 | 1.7 | 6.14 ± 0.03 | 6.5 ± 0.05 | 0.36 ± 0.06 | 2.3 |
| 4 | 7.59 ± 0.11 | 7.79 ± 0.09 | 0.2 ± 0.14 | 1.6 | 6.17 ± 0.04 | 6.85 ± 0.05 | 0.68 ± 0.06 | 4.8 |
| 5 | 7.62 ± 0.13 | 8.01 ± 0.06 | 0.39 ± 0.14 | 2.5 | 6.02 ± 0.05 | 6.48 ± 0.05 | 0.46 ± 0.07 | 2.9 |
| 6 | 7.44 ± 0.16 | 8.16 ± 0.07 | 0.72 ± 0.17 | 5.2 | 6.16 ± 0.04 | 6.99 ± 0.05 | 0.83 ± 0.06 | 6.8 |
| 7 | 7.58 ± 0.12 | 7.55 ± 0.05 | -0.03 ± 0.13 | 0.9 | 6.03 ± 0.05 | 6.81 ± 0.05 | 0.78 ± 0.07 | 6 |
| 8 | 7.59 ± 0.11 | 7.65 ± 0.11 | 0.06 ± 0.16 | 1.1 | 6.00 ± 0.04 | 6.72 ± 0.05 | 0.72 ± 0.06 | 5.2 |
| 9 | 7.49 ± 0.08 | 7.52 ± 0.16 | 0.03 ± 0.18 | 1.1 | 5.98 ± 0.05 | 6.49 ± 0.03 | 0.51 ± 0.06 | 3.2 |
| 10 | 7.61 ± 0.09 | 7.61 ± 0.1 | 0 ± 0.13 | 1 | 5.98 ± 0.06 | 6.35 ± 0.06 | 0.37 ± 0.08 | 2.3 |
| 11 | 7.27 ± 0.18 | 7.52 ± 0.1 | 0.25 ± 0.21 | 1.8 | 6.07 ± 0.05 | 6.75 ± 0.05 | 0.68 ± 0.07 | 4.8 |
| 12 | 7.74 ± 0.11 | 7.88 ± 0.09 | 0.14 ± 0.14 | 1.4 | 6.14 ± 0.04 | 6.53 ± 0.04 | 0.39 ± 0.06 | 2.5 |
| 13 | 7.61 ± 0.11 | 7.68 ± 0.05 | 0.07 ± 0.12 | 1.2 | 6.01 ± 0.03 | 6.33 ± 0.03 | 0.32 ± 0.04 | 2.1 |
| 14 | 7.85 ± 0.07 | 7.78 ± 0.09 | -0.07 ± 0.11 | 0.9 | 6.07 ± 0.03 | 6.28 ± 0.03 | 0.21 ± 0.04 | 1.6 |
| 15 | 7.71 ± 0.11 | 7.93 ± 0.05 | 0.22 ± 0.12 | 1.7 | 5.95 ± 0.06 | 5.95 ± 0.04 | 0 ± 0.07 | 1 |
| 16 | 7.94 ± 0.06 | 7.63 ± 0.1 | -0.31 ± 0.12 | 0.5 | 6.12 ± 0.02 | 6.55 ± 0.08 | 0.43 ± 0.08 | 2.7 |
| 17 | 7.67 ± 0.06 | 7.87 ± 0.04 | 0.2 ± 0.07 | 1.6 | 5.96 ± 0.06 | 6.28 ± 0.05 | 0.32 ± 0.08 | 2.1 |
| 18 | 7.67 ± 0.10 | 7.59 ± 0.12 | -0.08 ± 0.16 | 0.8 | 6.12 ± 0.04 | 6.88 ± 0.05 | 0.76 ± 0.06 | 5.8 |
| 19 | 7.68 ± 0.06 | 7.87 ± 0.06 | 0.19 ± 0.08 | 1.5 | 5.95 ± 0.04 | 6.35 ± 0.06 | 0.4 ± 0.07 | 2.5 |
| 20 | 7.92 ± 0.06 | 7.85 ± 0.06 | -0.07 ± 0.08 | 0.9 | 6.22 ± 0.04 | 6.66 ± 0.03 | 0.44 ± 0.05 | 2.8 |
| 21 | 6.80 ± 0.12 | 6.94 ± 0.14 | 0.14 ± 0.18 | 1.4 | 6.01 ± 0.05 | 6.79 ± 0.04 | 0.78 ± 0.06 | 6 |
| 22 | 7.73 ± 0.10 | 7.57 ± 0.13 | -0.16 ± 0.16 | 0.7 | 6.25 ± 0.05 | 6.53 ± 0.05 | 0.28 ± 0.07 | 1.9 |

**Table S5**: Allosteric parameters derived from $G\alpha oA$ BRET and β-arrestin2 BRET assays of methadone-, morphine-, oxycodone-, or DAMGO-bound MOR in the presence of increasing concentrations of Comp5 or MS1 (Compound #18).

| Opioid drugs | PAM | GαoA BRET | | | β-arrestin2 BRET | | | TTEST (p-value) | Bias |
|---|---|---|---|---|---|---|---|---|---|
| | | log(αβ) | $pK_B$ | log(αβ/$K_B$) | log(αβ) | $pK_B$ | log(αβ/$K_B$) | | |
| Methadone | Comp5 | 0.64 ± 0.14 | 6.03 | 6.67 ± 0.14 | 0.55 ± 0.24 | 6.03 | 6.58 ± 0.24 | 0.74 | Balanced |
| | MS1 | 0.85 ± 0.12 | 5.45 | 6.3 ± 0.12 | 0.79 ± 0.19 | 5.45 | 6.24 ± 0.19 | 0.81 | Balanced |
| Morphine | Comp5 | 0.51 ± 0.22 | 6.03 | 6.54 ± 0.22 | -0.02 ± 0.14 | 6.03 | 6.01 ± 0.14 | 0.04 | Modest G |
| | MS1 | 0.2 ± 0.21 | 5.45 | 5.65 ± 0.21 | 0.05 ± 0.17 | 5.45 | 5.5 ± 0.17 | 0.58 | Balanced |
| Oxycodone | Comp5 | 0.56 ± 0.24 | 6.03 | 6.59 ± 0.24 | 0.03 ± 0.1 | 6.03 | 6.06 ± 0.1 | 0.05 | Modest G |
| | MS1 | 0.2 ± 0.23 | 5.45 | 5.65 ± 0.23 | -0.03 ± 0.14 | 5.45 | 5.42 ± 0.14 | 0.4 | Balanced |
| DAMGO | Comp5 | 0.56 ± 0.14 | 6.03 | 6.59 ± 0.14 | 0.33 ± 0.19 | 6.03 | 6.36 ± 0.19 | 0.35 | Balanced |
| | MS1 | 0.53 ± 0.14 | 5.45 | 5.97 ± 0.14 | -0.06 ± 0.18 | 5.45 | 5.39 ± 0.18 | 0.01 | Modest G |

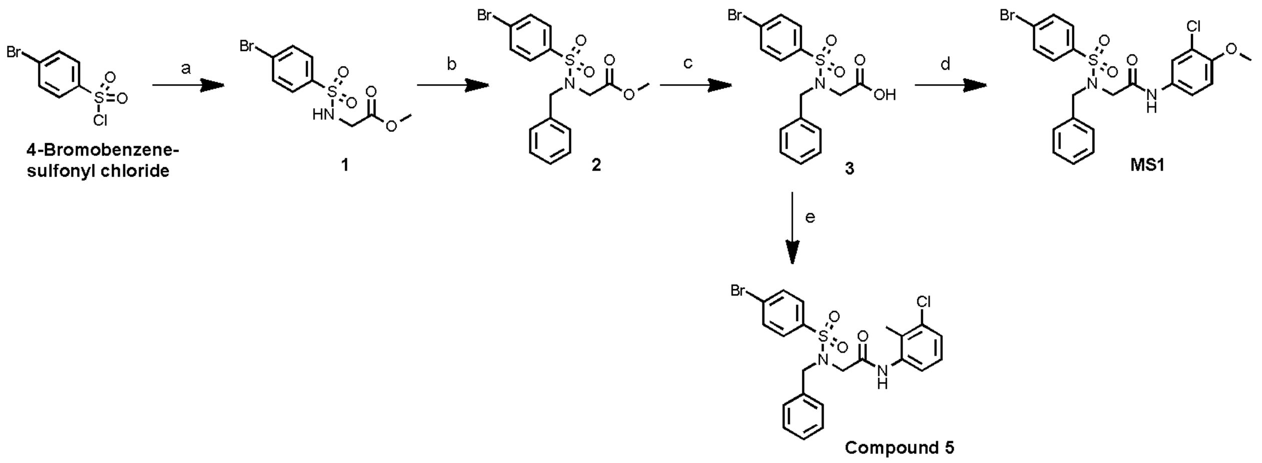


**Scheme 1. Synthetic Route, Reagents and Conditions for preparation of MS1 and Compound 5:** (a) glycine methyl ester hydrochloride, triethylamine, dichloromethane (DCM), 0 °C to room temperature (rt), 67%; (b) sodium hydride, benzyl bromide, dimethylformamide, 0 °C to rt, 74%; (c) lithium hydroxide, tetrahydrofuran, methanol, $H_2O$, rt, overnight, 85%; (d) 3-Chloro-4-methoxyaniline, 1-(3-Dimethylaminopropyl)-3-ethylcarbodiimide hydrochloride (EDCI), 1-Hydroxy-7-azabenzotriazole (HOAt), N,N-Diisopropylethylamine (DIEA), DCM, rt, overnight, 86%. (e) 3-Chloro-2-methylaniline, 1-(3-Dimethylaminopropyl)-3-ethylcarbodiimide hydrochloride (EDCI), 1-Hydroxy-7-azabenzotriazole (HOAt), N,N-Diisopropylethylamine (DIEA), DCM, rt, overnight, 76%.

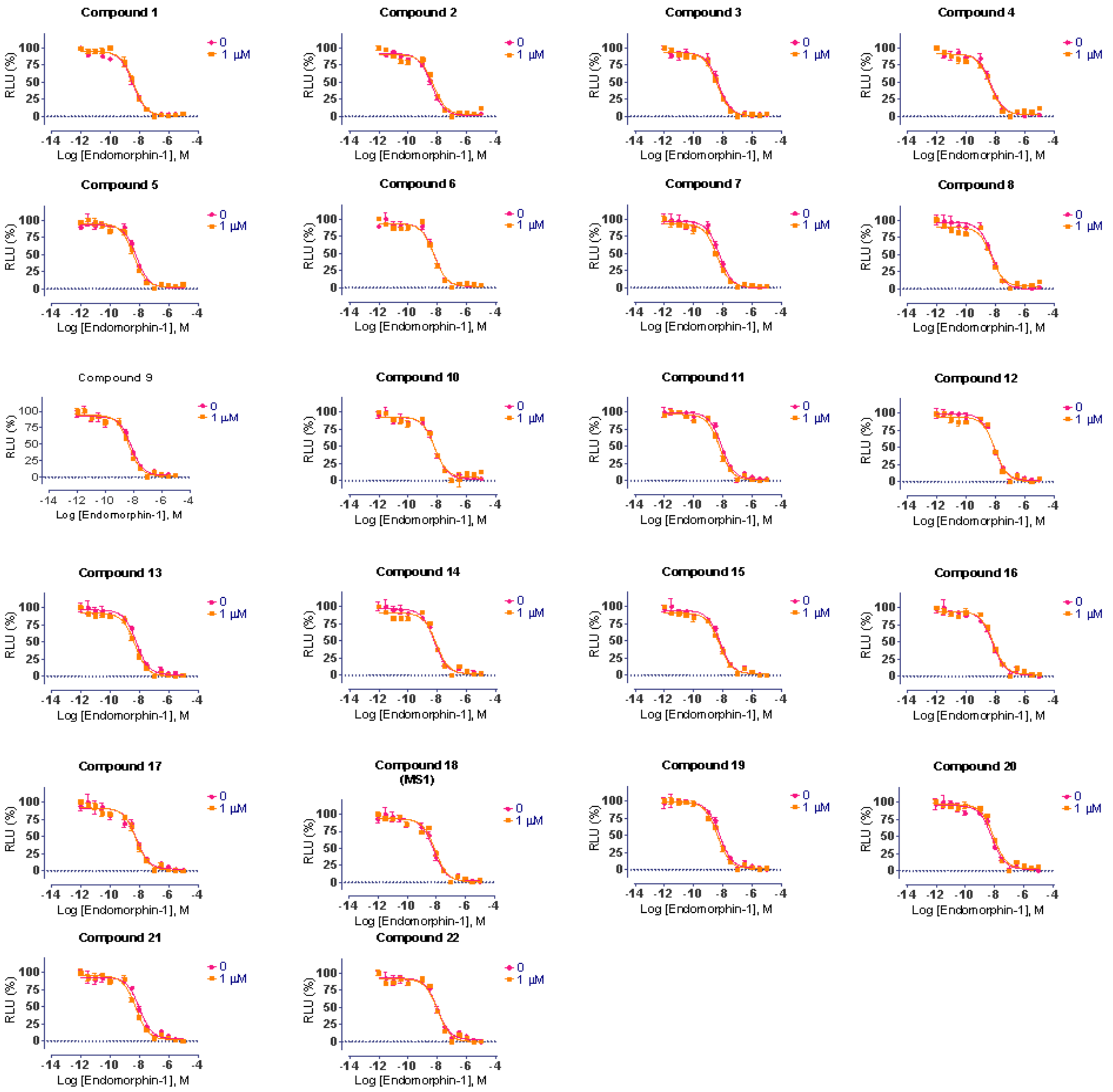


**Fig. S1. Concentration-response curves with graded concentrations of MS1 (compound #18) and derivatives using endomorphin-1 as an orthosteric agonist in cAMP assays measuring G protein activation.** Curves drawn in magenta and orange represent data at 0 µM (in the absence of a compound) and 1 µM compound concentrations, respectively.

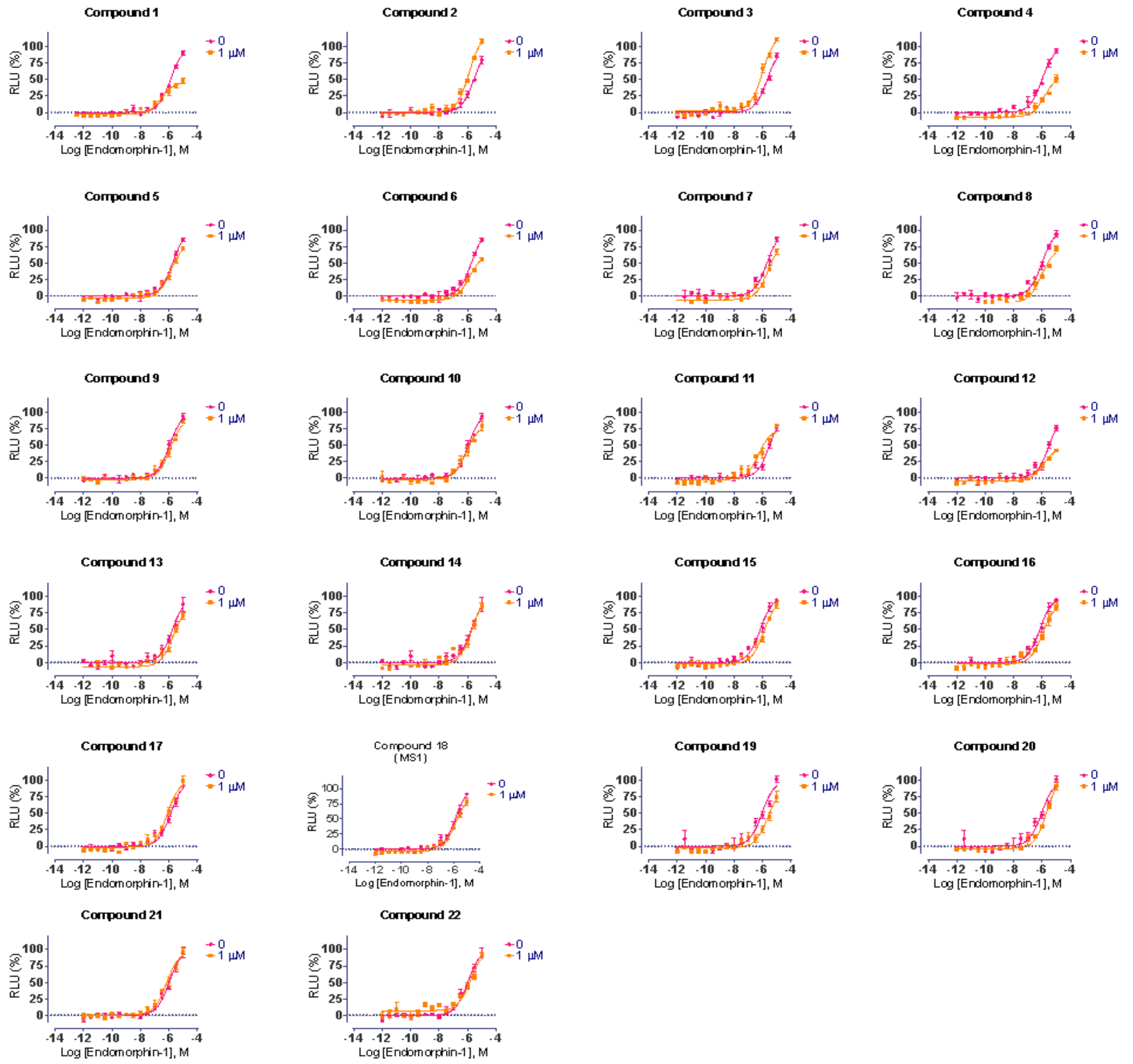


**Fig. S2. Concentration-response curves with graded concentrations of MS1 (compound #18) and derivatives using endomorphin-1 as an orthosteric agonist in Tango assays measuring β-arrestin recruitment.** Curves drawn in magenta and orange represent data at 0 µM (in the absence of a compound) and 1 µM compound concentrations, respectively.

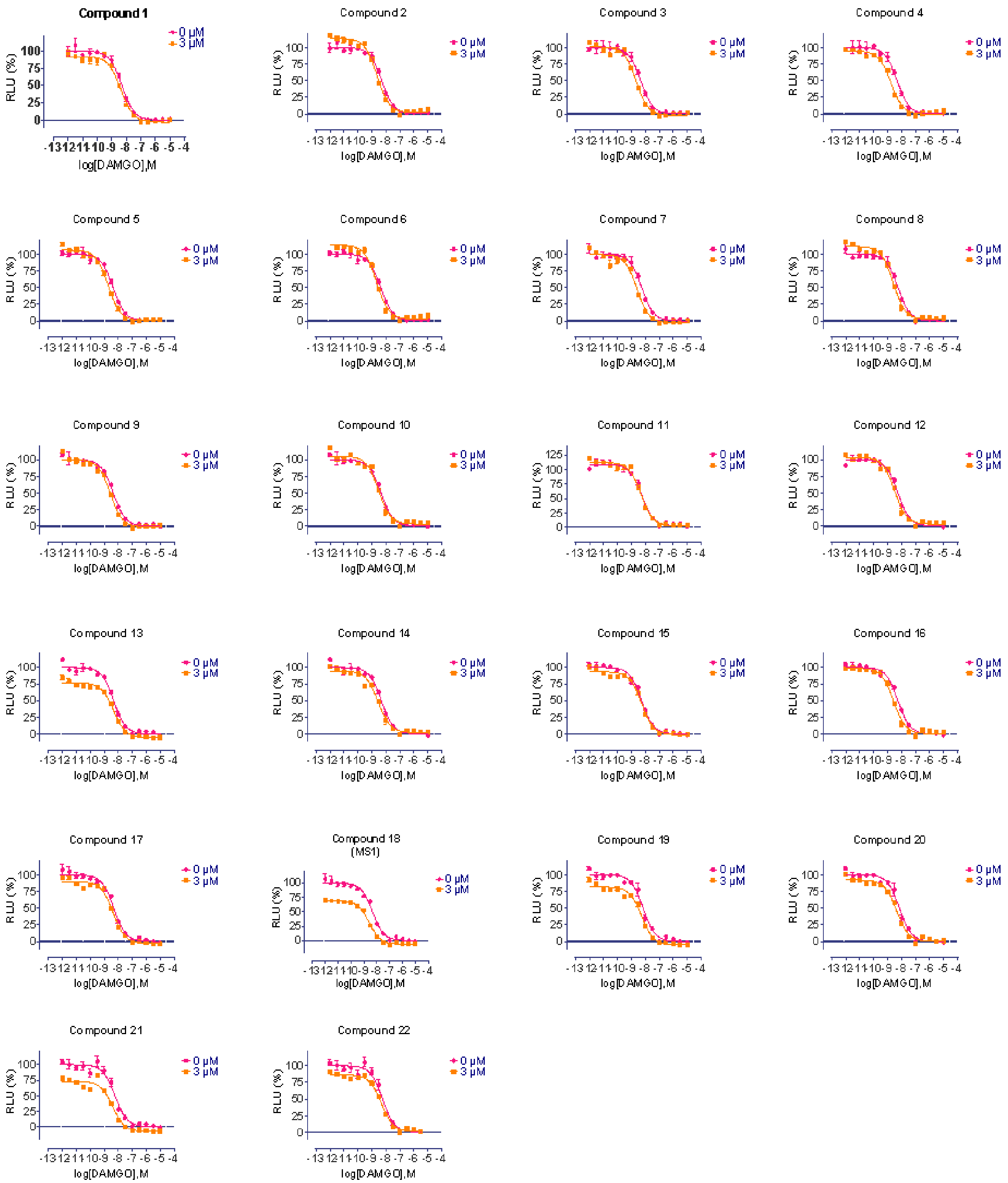


**Fig. S3. Concentration-response curves with graded concentrations of MS1 (compound #18) and derivatives using DAMGO as an orthosteric agonist in cAMP assays measuring G protein activation.** Curves drawn in magenta and orange represent data at 0 µM (in the absence of a compound) and 3 µM compound concentrations, respectively.

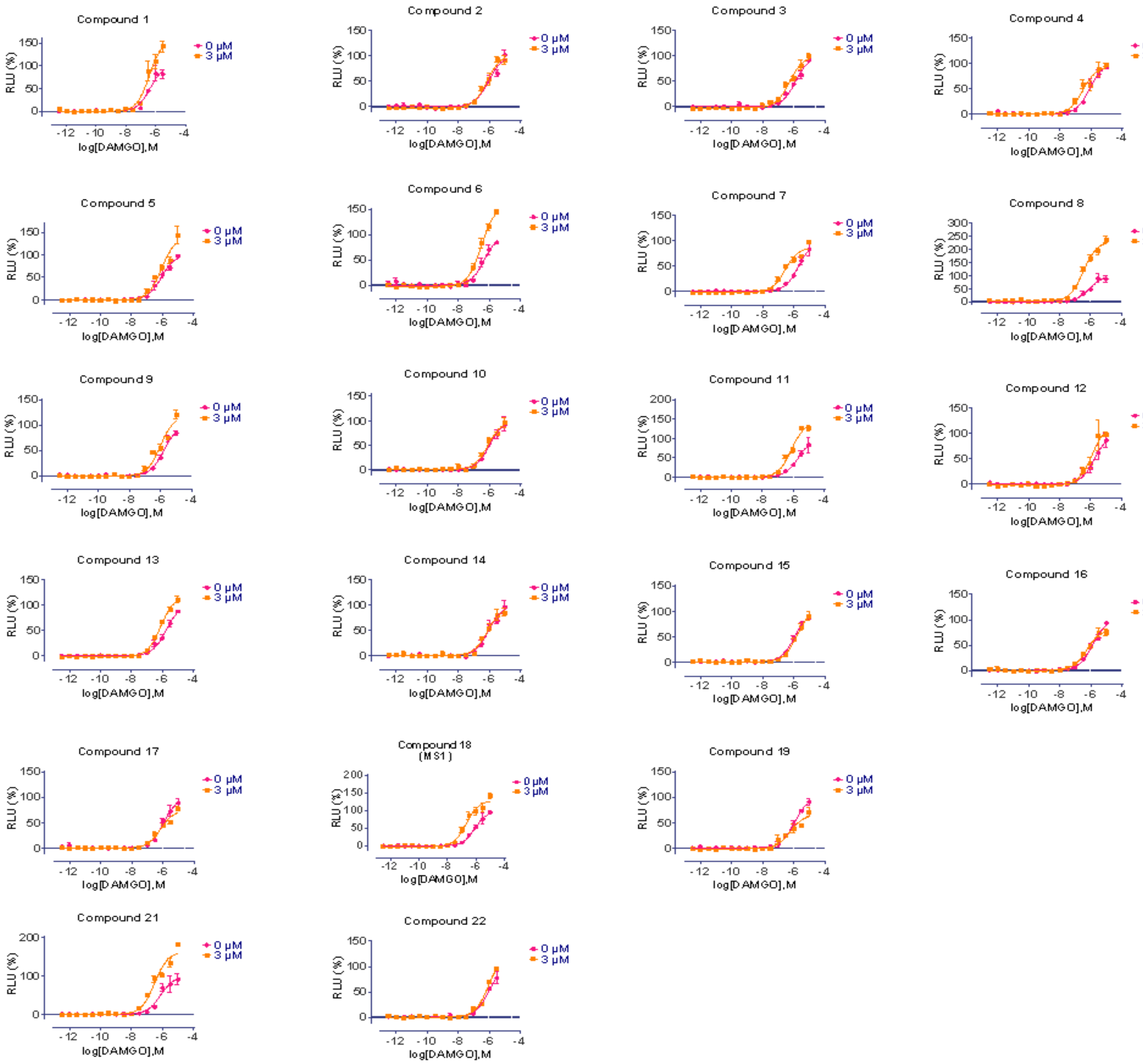


**Fig. S4. Concentration-response curves with graded concentrations of MS1 (compound #18) and derivatives using DAMGO as an orthosteric agonist in Tango assays measuring β-arrestin recruitment.** Curves drawn in magenta and orange represent data at 0 µM (in the absence of a compound) and 3 µM compound concentrations, respectively.

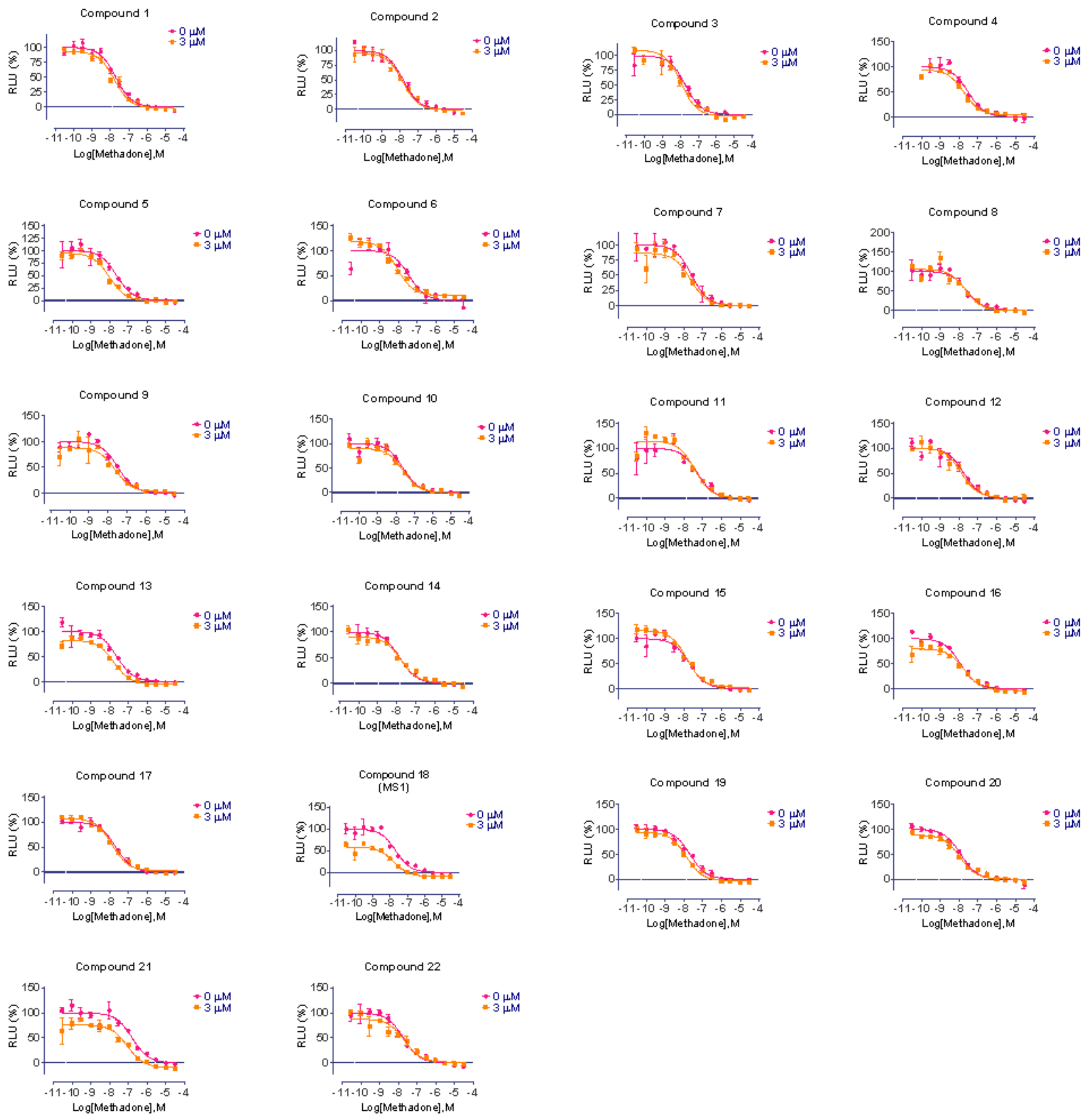


**Fig. S5. Concentration-response curves with graded concentrations of MS1 (compound #18) and derivatives using methadone as an orthosteric agonist in cAMP assays measuring G protein activation.** Curves drawn in magenta and orange represent data at 0 µM (in the absence of a compound) and 3 µM compound concentrations, respectively.

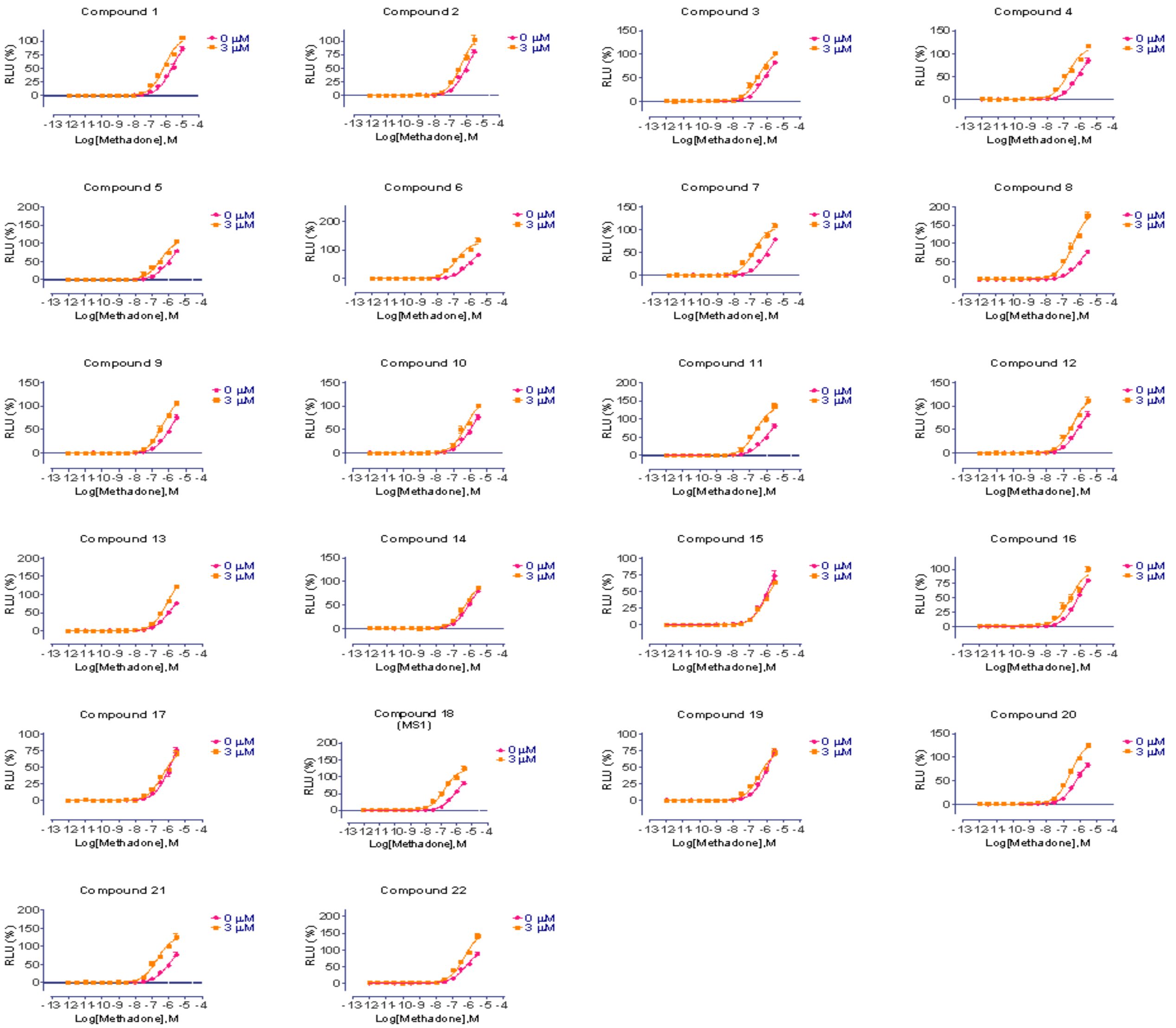


**Fig. S6. Concentration-response curves with graded concentrations of MS1 (compound #18) and derivatives using methadone as an orthosteric agonist in Tango assays measuring β-arrestin recruitment.** Curves drawn in magenta and orange represent data at 0 µM (in the absence of a compound) and 3 µM compound concentrations, respectively.

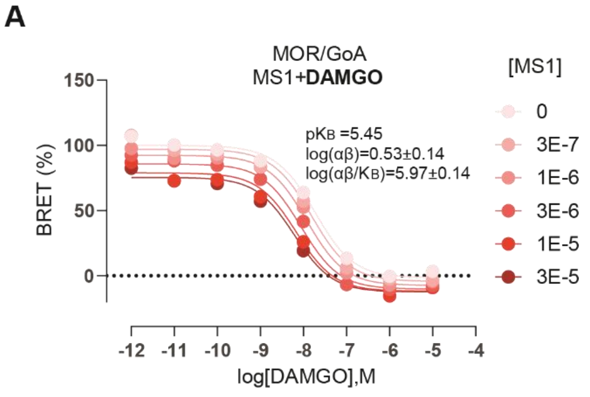


| | 0 | 3E-7 | 1E-6 | 3E-6 | 1E-5 | 3E-5 |
|---|---|---|---|---|---|---|
| LogEC50 | -7.787 | -7.863 | -7.851 | -7.992 | -8.173 | -8.346 |

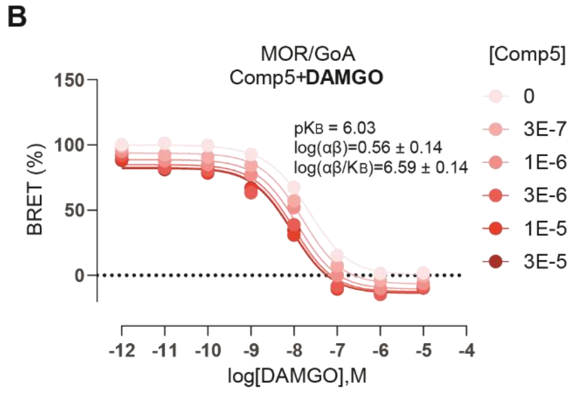


| | 0 | 3E-7 | 1E-6 | 3E-6 | 1E-5 | 3E-5 |
|---|---|---|---|---|---|---|
| LogEC50 | -7.707 | -7.806 | -7.814 | -8.007 | -8.128 | -8.066 |

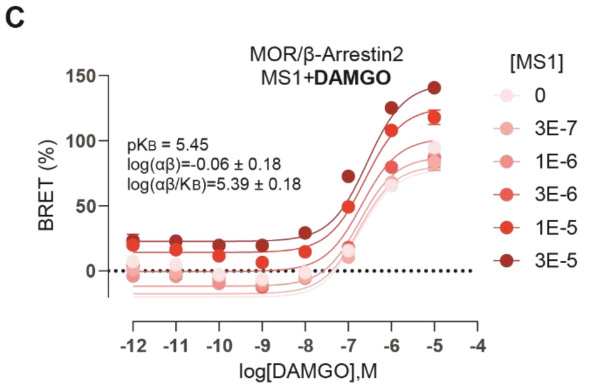


| | 0 | 3E-7 | 1E-6 | 3E-6 | 1E-5 | 3E-5 |
|---|---|---|---|---|---|---|
| LogEC50 | -6.274 | -6.414 | -6.515 | -6.582 | -6.715 | -6.865 |

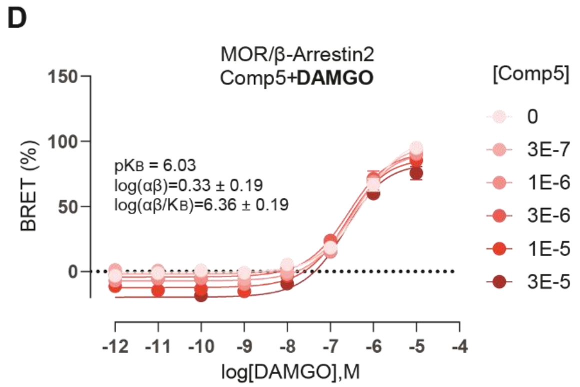


| | 0 | 3E-7 | 1E-6 | 3E-6 | 1E-5 | 3E-5 |
|---|---|---|---|---|---|---|
| LogEC50 | -6.331 | -6.360 | -6.462 | -6.601 | -6.748 | -6.746 |

**Fig. S7. Concentration-response curves of MOR with graded concentrations of MS1 and Comp5 using DAMGO as the orthosteric ligand.** GαoA dissociation BRET assay of (A) MS1 and (B) Comp5 and β-arrestin recruitment BRET assays of (C) MS1 and (D) Comp5 were performed to assess MS1 and Comp5 allosteric regulation of DAMGO-induced signaling through MOR. Data are mean +/- SEM of a dataset composed of three independent experiments, each one carried out in duplicate. $K_B$ is the binding affinity of PAM to free receptor, α is the binding cooperativity factor, and β is the efficacy cooperativity factor.

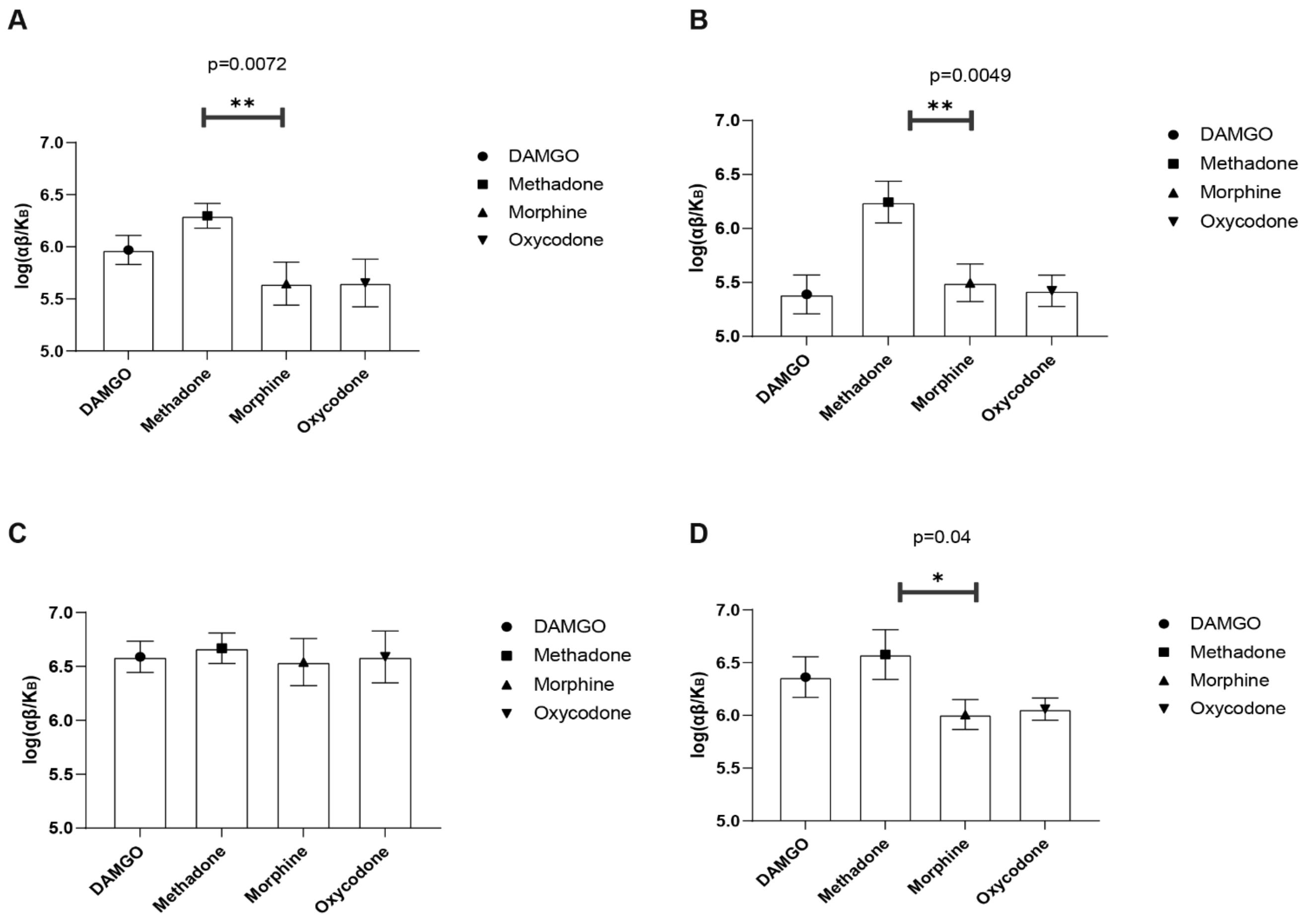


**Fig. S8. Different probe dependence of MS1 and Comp5.** The $\log(\alpha\beta/K_B)$ allosteric modulation estimates of (A) MS1 in GαoA BRET assay, (B) MS1 in β-arrestin recruitment BRET assay, (C) Comp5 in GαoA dissociation BRET assay and (D) Comp5 in β-arrestin recruitment BRET assays calculated for different opioid drugs and represented as bar graphs. Data are mean +/- SEM of a dataset composed of three independent experiments, each one carried out in duplicate. $K_B$ is the binding affinity of PAM to free receptor, α is the binding cooperativity factor, and β is the efficacy cooperativity factor. The statistical significance of $\log(\alpha\beta/K_B)$ differences between methadone and morphine was assessed by t-test.

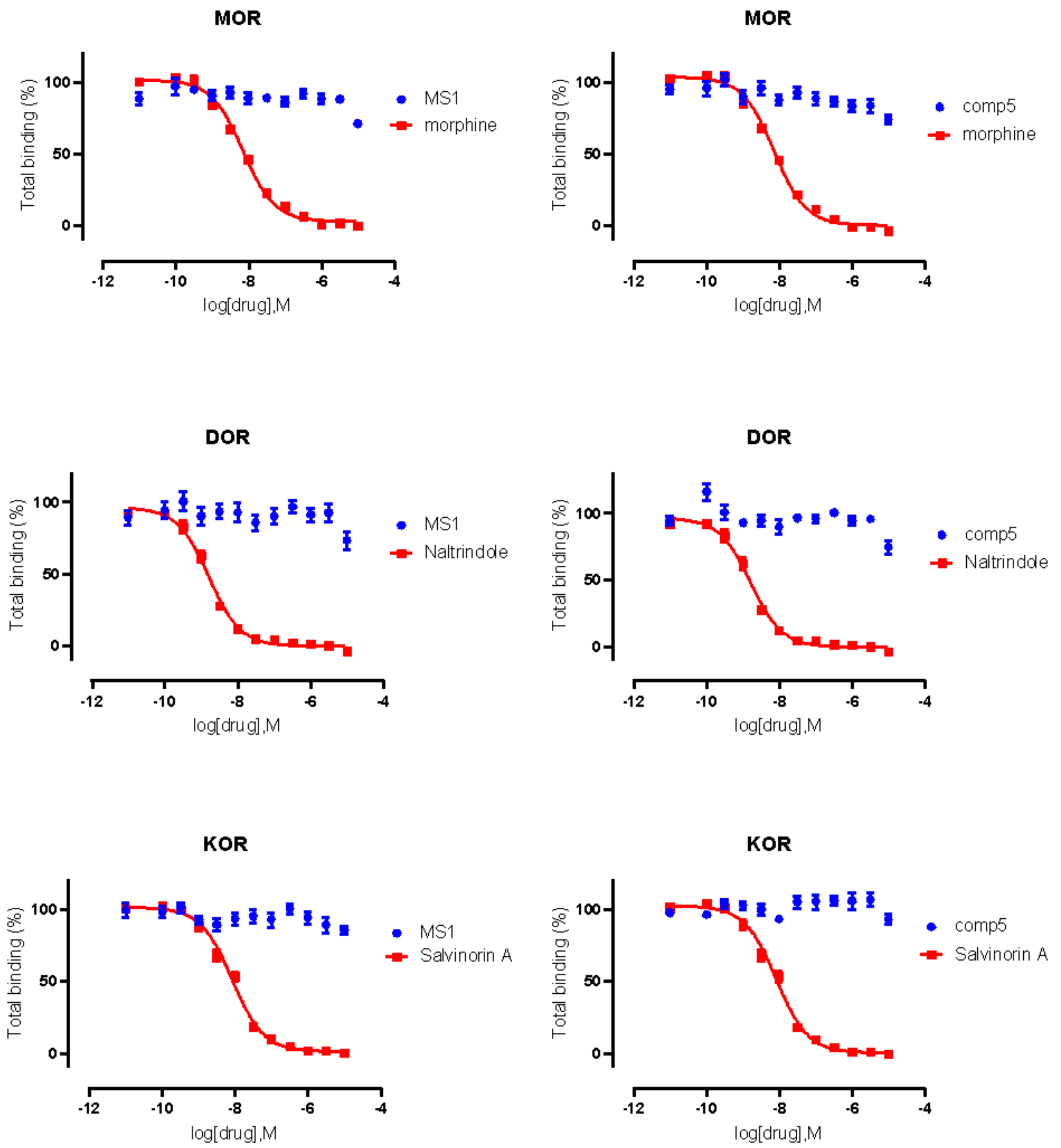


**Fig. S9. The binding affinity of Comp5 and MS1 on human MOR, DOR or KOR.** [$^{3}$H]-DAMGO, [$^{3}$H]-DADLE, and [$^{3}$H]-U69593 were used as radioligands for MOR, DOR, and KOR, respectively. Reference ligands were also tested in parallel with MS1 and Comp5.

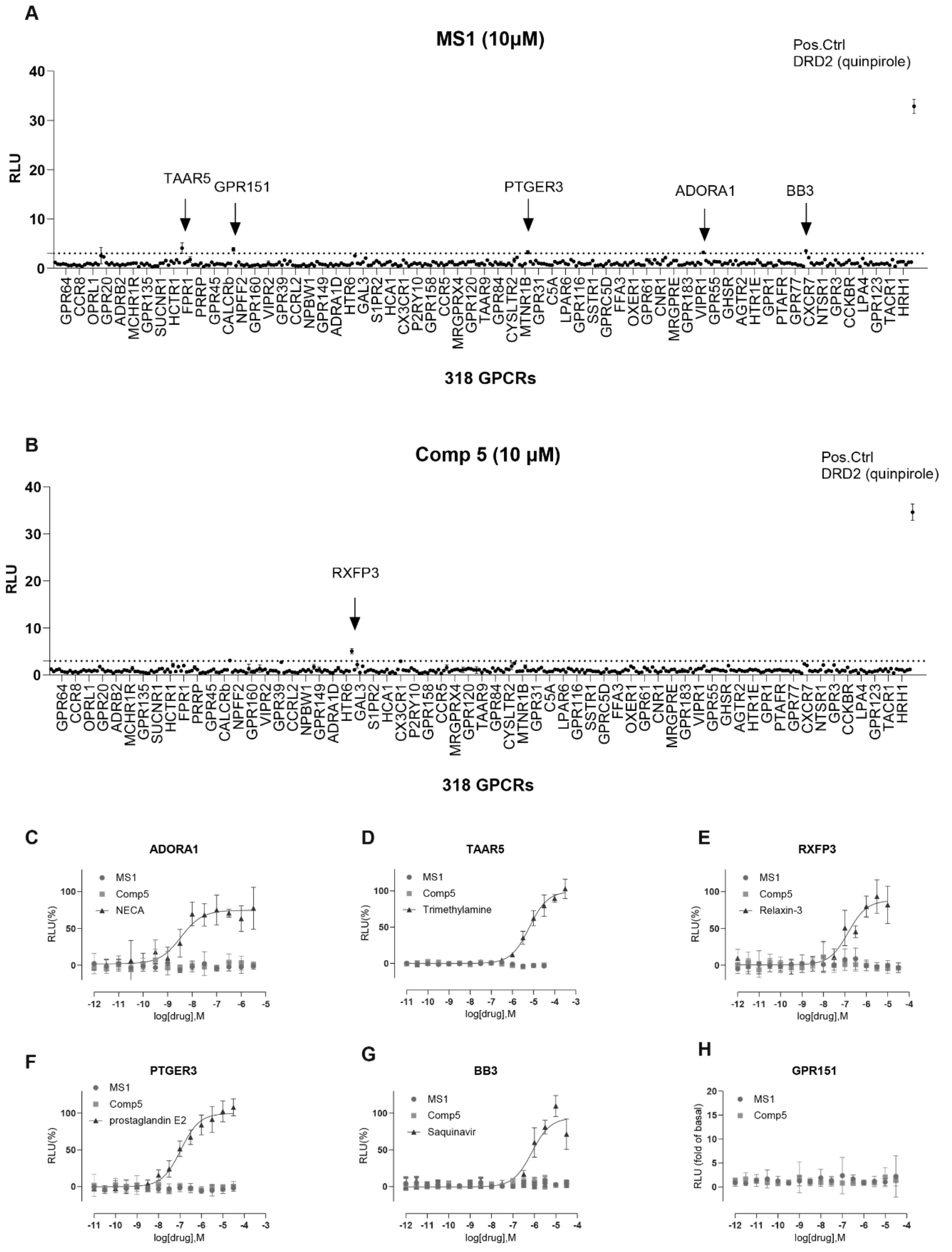


**Fig. S10. Screening of MS1 and Comp5 in single concentrations (10 μM) at 318 GPCRs** in PRESTO-Tango GPCRome assay (only ~20% receptors are shown in panels A and B, respectively) and follow-up dose response curves (panels C-H) for those receptors that showed activity (RLU>3). DRD2 (quinpirole) in panels A and B indicates the known DRD2 agonist quinpirole, which is used as a positive control to show that the system is working. Panels C-H show dose response curves of MS1, Comp5, and known agonists (positive controls) for each receptor that showed activity, except GPR151, which does not have a reference ligand (results are plotted as fold of basal in this case).

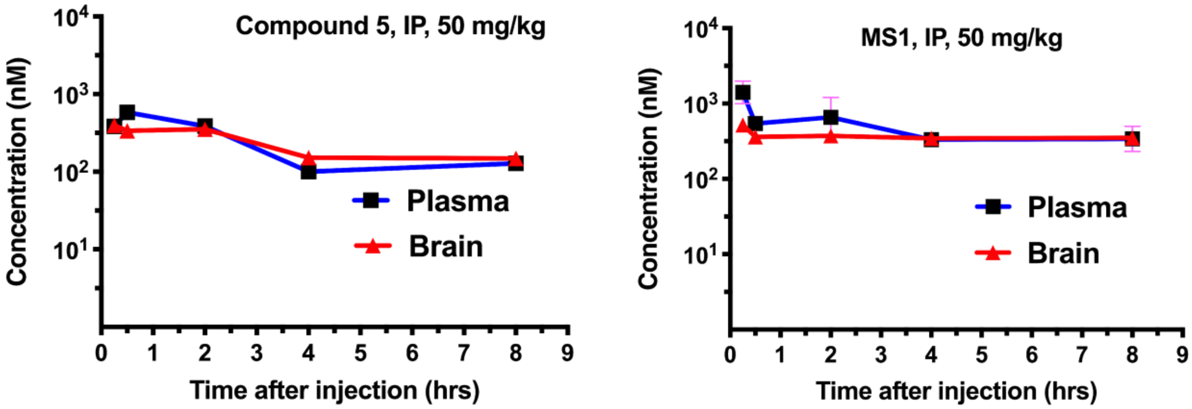


**Fig. S11. Comp5 and MS1 brain and plasma concentrations** after a single 50 mg/kg i.p. injection in mice (average values, 3 mice/time point).

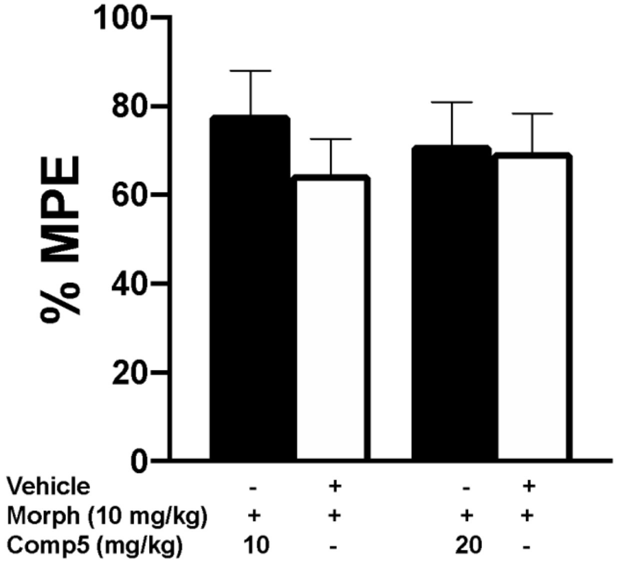


**Fig. S12. Low doses of Comp5 do not affect the antinociceptive actions of morphine in the hot plate assay.** Male mice were injected with Comp5 or vehicle immediately after hot plate baseline assessment, received either opioid or saline treatment 30 min later, and they were tested in the hot plate apparatus 30 min after the last injection. Data are plotted as % maximal possible effect (MPE). Comp5 at doses of 10 and 20 mg/kg does not have any effect on the antinociceptive efficacy of morphine (n=7-8 per group). Data are reported as ± SEM. Statistics based on multiple comparisons one-way ANOVA followed by Holm—Sidak's post hoc test. Morph (Morphine) and Comp5 (Compound-5).Comp5 was injected i.p. and morphine was injected s.c.

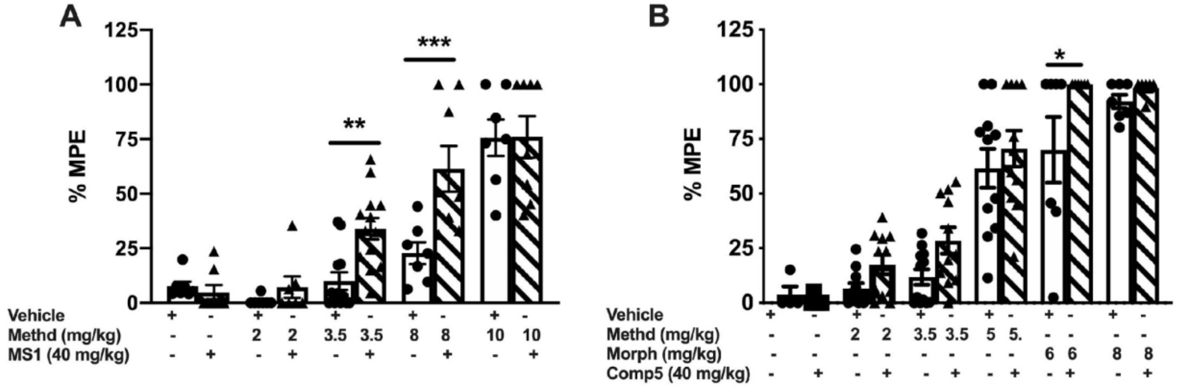


**Fig. S13. MS1 potentiates the analgesic actions of methadone and Comp5 potentiates the effects of morphine in female mice in the hot plate assay.** Female mice were injected with MS1, Comp5 or vehicle immediately after hotplate baseline assessment, received either methadone or saline treatment 30 min later, and were tested 30 min later. Data are plotted as % maximal possible effect (MPE). (A) MS1 increases the efficacy of methadone [$F(9,74)= 22.62$, $p<0.001$, (n=7-13 per group), **$p<0.01$ for methadone 3.5 mg/kg + Veh vs. methadone 3.5 mg/kg + MS1, ***$p<0.001$ for methadone 8 mg/kg vs. methadone 8 mg/kg + compound-] and (B) Comp5 has no effect on the analgesic efficacy of methadone (n=5-12) but enhances the analgesic efficacy of morphine [$F(11,90)= 28.83$, $p<0.0001$, (n=6-7), **$p<0.05$ for morphine 6 mg/kg +Veh vs. morphine 6 mg/kg + Comp5. Data are reported as ± SEM. Statistics based on multiple comparisons one-way ANOVA followed by Holm—Sidak's post hoc test. Morph (Morphine), Methd (Methadone) and Comp5 (Compound-5). MS1, Comp5, or vehicle were injected i.p. and all opioids were injected s.c.

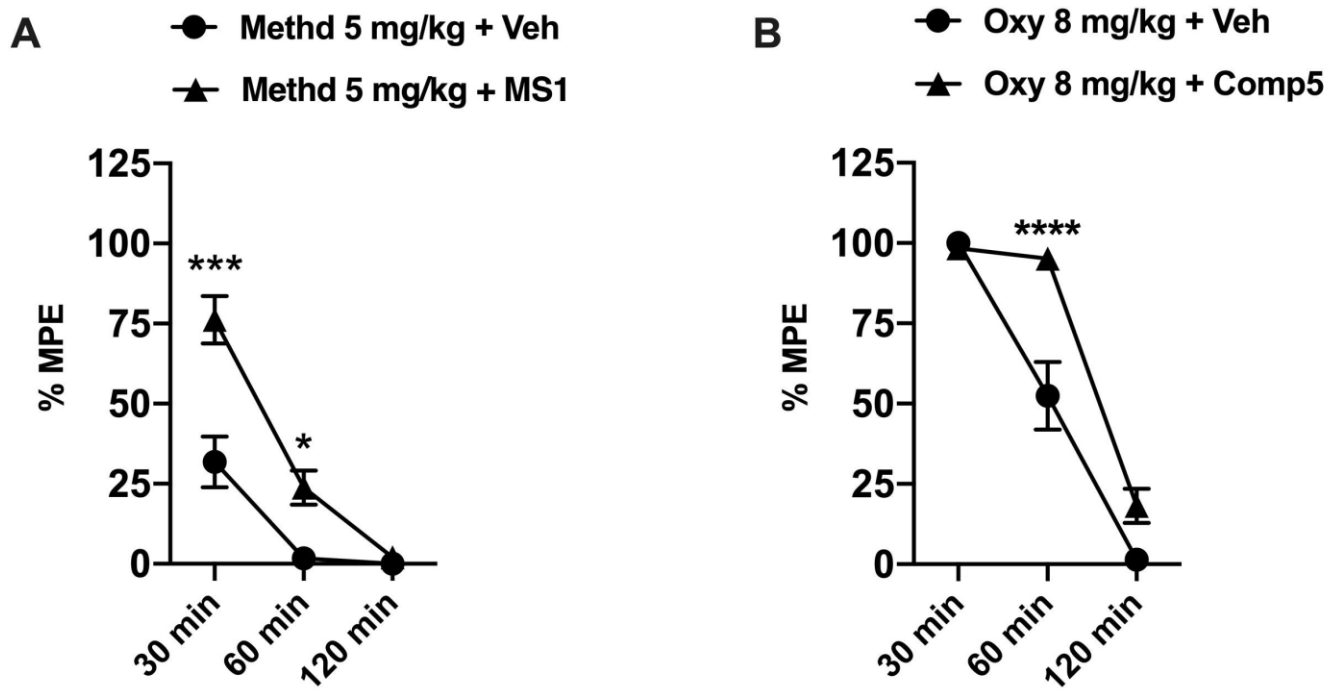


**Fig. S14. Pretreatment with MS1 and Comp5 prolong the antinociceptive effects of methadone and oxycodone in the hot plate assay.** Male mice were injected with Comp5, MS1, or vehicle immediately after hotplate baseline assessment, received opioid treatment 30 min later, and were tested at 30 min, 60 min, and 120 min after the opioid injection. Data are plotted as % maximal possible effect (MPE). (A) MS1 enhances the duration of analgesic effect of methadone in the hot plate assay [A significant interaction was reported between treatment x time $F(2,10) = 11.87$, $p=0.0023$, (n=7),***$p<0.001$ for Methadone 5 mg/kg _+ Veh vs. Methadone 5 mg/kg + MS1 at 30 min, *$p<0.05$ for Methadone 5 mg/kg + Veh vs. Methadone 5 mg/kg + MS1 at 60 min]. (B) Comp5 enhances the duration of analgesic effect of oxycodone in the hot plate assay [A significant interaction was reported between treatment x time $F(2,14) = 12.42$, $p=0.0008$, (n=8),***$p<0.0001$ for Oxycodone 8 mg/kg + Veh vs. Oxycodone 8 mg/kg + Comp5 at 60 min]. Data are reported as ± SEM. Statistics based on multiple comparisons two-way ANOVA followed by Holm—Sidak's post hoc test. Methd (Methadone), Veh (Vehicle), and Comp5 (Compound-5). MS1, Comp5, or vehicle were injected i.p. and all opioids were injected s.c.

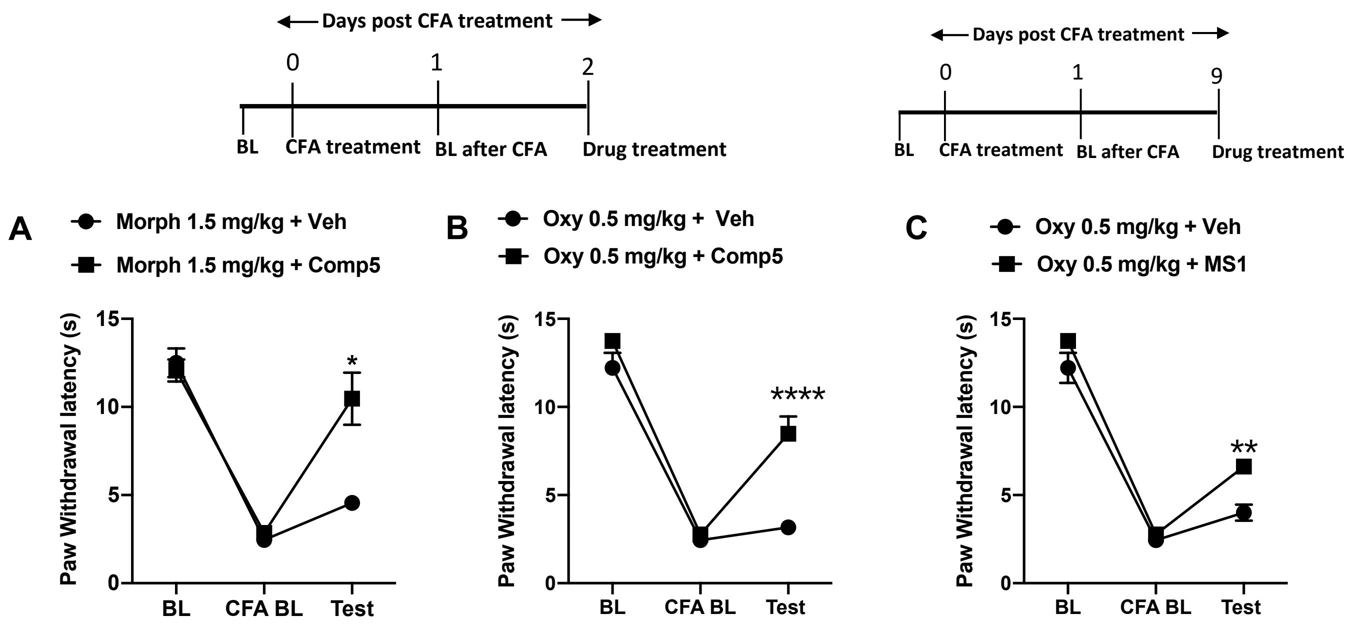


**Fig. S15**. **PAMs promote the antihyperalgesic effects of opioids in models of peripheral inflammation in female mice.** CFA treatment was administered to the left hindpaw after taking baseline (BL) measurements. Baseline CFA (CFA BL) measurement was taken 24 hours after CFA treatment. Female mice were injected with Comp5, MS1 or vehicle then received opioid treatment 30 min later, and Hargreaves responses were tested 30 min after the opioid injection. Data are reported as paw withdrawal latency in seconds. (A) Comp5 promotes antihyperalgesic responses to morphine [A significant interaction was reported between treatment x test time F (2,26)= 9.345, p=0.0009, (n=7-8 per group), *p<0.05 for morphine 1.5 mg/kg +Veh vs. morphine 1.5 mg/kg + Comp5]. (B) Comp5 promotes antihyperalgesic responses to oxycodone. [A significant interaction was reported between treatment x test time F (2,26)= 8.403, p= 0.0015, (n=7-8 per group), ****p<0.001 for oxycodone 0.5 mg/kg +Veh vs. oxycodone 0.5 mg/kg + Comp5]. (C) MS1 promotes antihyperalgesic responses to oxycodone [The interaction between treatment x time F(2,26)= 2.915, p=0.0721, (n=7-8), **p<0.01 for oxycodone 0.5 mg/kg vs. oxycodone 0.5 mg/kg + MS1]. Data are reported as ± SEM. Statistics based on multiple comparisons two-way ANOVA followed by Holm—Sidak's post hoc test (A and B) and Bonferroni post hoc test (C). Morph (Morphine), Oxy (Oxycodone), and Comp5 (Compound-5). MS1, Comp5, or vehicle were injected i.p. and all opioids were injected s.c.

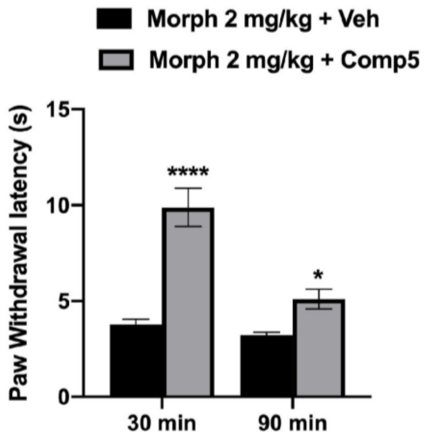


**Fig. S16. Comp5 produced a small but significant increase in the duration of morphine antinociception.** Time course experiments were conducted with the same group of mice used in Fig.3A and an additional measurement was taken 90 min after the morphine injection [A significant interaction was reported between treatment x time $F(1,14)= 26.33$, $p=0.0002$, (n=8), ****$p<0.0001$ for morphine 2 mg/kg vs. morphine 2 mg/kg + Comp5 at 30 min, *$p=0.0299$ for morphine 2 mg/kg vs. morphine 2 mg/kg + Comp5 at 90 min]. Morph (Morphine) and Comp5 (Compound-5). Comp5 was injected i.p. and morphine was injected s.c.
